# Plant geographic distribution influences chemical defenses in native and introduced *Plantago lanceolata* populations

**DOI:** 10.1101/2023.06.05.543708

**Authors:** Pamela Medina-van Berkum, Eric Schmöckel, Armin Bischoff, Natalia Carrasco-Farias, Jane A. Catford, Reinart Feldmann, Karin Groten, Hugh A. L. Henry, Anna Lampei Bucharova, Sabine Hänniger, Justin C. Luong, Julia Meis, Vincensius S.P. Oetama, Meelis Pärtel, Sally A. Power, Jesus Villellas, Erik Welk, Astrid Wingler, Beate Rothe, Jonathan Gershenzon, Michael Reichelt, Christiane Roscher, Sybille B. Unsicker

**Author notes:** ***Corresponding authors:*** Sybille B. Unsicker, Christiane Roscher. Denotes equal authorship contribution (Pamela Medina-van Berkum, Eric Schmöckel). **Author’s contributions** CR and SBU designed the study; AB, NCF, JC, RF, HALH, ALB, JCL, JM, MNG, MP, SAP, JV and AW collected the seed material; SH and VSPO provided the caterpillars for this study, KG organized the collection and importation of the seeds. EW compiled the distribution data set; ES, PMB, BR and MR performed the experiment and chemical analysis. PMB and ES analyzed the data; PMB, ES, SBU, CR and JG wrote the first draft of the manuscript. All co-authors discussed the results, contributed substantially to the drafts, and gave final approval of the manuscript prior to the submission.

## Abstract

Plants growing outside their native range may be confronted by new regimes of herbivory, but how this affects plant chemical defense profiles has rarely been studied. Using *Plantago lanceolata* as a model species, we investigated whether introduced populations show significant differences from native populations in several growth and chemical defense traits. *Plantago lanceolata* (ribwort plantain) is an herbaceous plant species native to Europe and Western Asia that has been introduced to numerous countries worldwide. We sampled seeds from nine native and ten introduced populations that covered a broad geographic and environmental range and performed a common garden experiment in a greenhouse, in which we infested half of the plants in each population with caterpillars of the generalist herbivore *Spodoptera littoralis*. We then measured size-related and resource-allocation traits as well as the levels of constitutive and induced chemical defense compounds in roots and shoots of *P. lanceolata*. When we considered the environmental characteristics of the site of origin, our results revealed that populations from introduced ranges were characterized by an increase of chemical defense compounds without compromising plant biomass. The concentrations of iridoid glycosides and verbascoside, the major anti-herbivore defense compounds of *P. lanceolata*, were higher in introduced populations than in native populations. In addition, introduced populations exhibited greater rates of herbivore-induced volatile organic compound emission and diversity, and similar chemical diversity based on untargeted analyses of leaf methanol extracts. In general, the geographic origin of the populations had a significant influence on morphological and chemical plant traits, suggesting that *P. lanceolata* populations are not only adapted to different environments in their native range, but also in their introduced range.

## Introduction

The range of many plant species has been extended into new regions by accidental or deliberate anthropogenic introduction (Mack & Lonsdale 2001). Plants may exhibit new phenotypes in response to changing biotic and abiotic environmental factors in their new range (van Kleunen, Bossdorf & Dawson 2018). Studies comparing native and introduced plant populations often emphasize the importance of chemical defenses against herbivores because they are considered critical in controlling plant establishment, development, and survival (Cipollini et al. 2005; Huang et al. 2010; Wolf et al. 2011). However, most of these studies have examined only a few plant populations and a small fraction of defense metabolites. Therefore, it remains unclear whether the establishment of a plant species in a new range leads to a rapid evolutionary change in the composition of its defenses.

Various non-mutually exclusive hypotheses have been proposed to explain the mechanisms underlying the establishment and dominance of some introduced species. Introduced plants may experience reduced biotic selection pressures due to partial or complete release from pathogens and herbivores found in their native range (ERH: Enemy release hypothesis; Keane & Crawley 2002). They can thus reallocate resources from defense to growth or reproduction, leading to enhanced competitiveness (EICA: Evolution of increased competitive ability hypothesis; Blossey & Nötzold 1995; Bossdorf *et al*. 2005), or shift defense strategies towards less-costly chemical compounds (SDH: Shifting defense hypothesis). Moreover, under some conditions, they might also benefit from the synthesis of new defense compounds (NWH: Novel weapon hypothesis; Callaway & Ridenour 2004). Numerous studies have tested these hypotheses with varying degrees of support (Yannelli et al. 2020) suggesting that the response and successful establishment of a plant species in a new range might vary depending on genetic diversity, spatial-temporal scales, and environmental conditions (Theoharides & Dukes 2007; Bock *et al*. 2015; Smith *et al*. 2020; Catford *et al*. 2021).

The introduction of plants into new ranges often occurs over a large area, spanning different countries and continents, such that new populations are exposed to a wide variety of different climatic conditions, soil properties, and biotic factors. As a result, introduced populations may show local adaptations (Hunter 2016). Differing intensities of biotic and abiotic factors might lead to clinal patterns in plant chemical profiles (Moreira *et al*. 2018). It has been hypothesized that investment in plant defense decreases at higher latitudes and altitudes due to lower herbivory pressure and diversity than in tropical environments or at lower elevations (Latitudinal / elevational herbivory defense hypothesis, Coley & Aide 1991; Pellissier *et al*. 2014). Although many studies provide explicit support for this hypothesis (e.g. Rasmann & Agrawal 2011), others do not and suggest that many plant-herbivore interactions are too variable to result in such a uniform pattern (Cremieux *et al*. 2008; Moles *et al*. 2011; Anstett et al. 2015).

Recent studies have found evidence that the chemical profiles of introduced populations differ from those of populations in their native range. For example, introduced individuals of *Tanacetum vulgare* L. showed increased concentrations of terpenes and other volatile compounds (Wolf *et al*. 2011). A study of native and introduced populations of *Ambrosia artemisiifolia* L. showed that phenolic compound composition did not only vary between ranges but also with latitude (van Boheemen *et al*. 2019). Similarly, the defense chemistry of *Phragmites australis* (Cav.) Trin. ex Steud. was strongly influenced by biogeographic range and latitude. Native genotypes had higher total phenolic contents than introduced genotypes, and concentrations of phenolics increased with latitude in native populations but not in introduced populations (Bhattarai et al. 2017).

In this study, we used *Plantago lanceolata* L. (ribwort plantain) to investigate how the colonization of a new range may lead to changes in chemical defense compounds. *Plantago lanceolata* is a common forb native to Europe and Western Asia that has been introduced and successfully established in numerous countries worldwide (Alexander *et al*. 2012; Penczykowkski & Sieg 2021), with records in North America, New Zealand, Australia and Japan dating back 200 years (Cavers, Bassett & Crompton 1980). Aside from a high degree of neutral and adaptive genetic diversity in both introduced and native ranges (Smith *et al*. 2020), *P. lanceolata* displays plastic and adaptive responses to environmental variation (Bischoff *et al*. 2006; Skinner & Stewart 2014). Different populations of *P. lanceolata* exhibit different metabolic profiles (Iwanycki Ahlstrand *et al*. 2018) but there is relatively little knowledge on how herbivore-induced metabolite patterns vary in this plant species across a geographic gradient. There are also numerous studies on the chemical defenses of *Plantago lanceolata*. For example, the iridoid glycosides (e.g. aucubin and catalpol) and phenylethanoid glycosides (e.g. verbascoside) are characteristic defense compounds against insect herbivores and pathogens. Their concentrations can be increased by insect herbivory (Darrow & Bowers 1999; Marak, Biere & Van Damme 2003), but also change in response to light, nutrient availability, mycorrhizal status, and neighboring plant species composition (Fontana *et al*. 2009; Mraja *et al*. 2011; Miehe-Steier *et al*. 2015).

To examine whether *P. lanceolata* populations change after their establishment in new geographic ranges, we quantified levels of chemical defense compounds in nine native and ten introduced populations that covered a broad range of geographic and environmental conditions. We asked the following questions: 1) how do native and introduced populations differ in size-related and resource-acquisition traits for the native and the introduced populations? 2) What are the differences in chemical traits both prior to and in response to insect herbivory? 3) Does the local climate of native and introduced populations influence their levels of chemical defense?

To address these questions, we performed a greenhouse experiment in which we infested half of the plants in each experimental population with generalist *Spodoptera littoralis* caterpillars. Using GC-MS and HPLC-MS, we compared constitutive and induced chemical defense compounds, both volatile and non-volatile from native and introduced populations of *P. lanceolata* and related them to geographic and climatic variables in the area of plant origin. We expected that plants from introduced ranges would have lower concentrations of defense compounds and higher biomass, consistent with the ERH, EICA and SDH hypotheses. Our results however, show that introduced populations were better defended chemically in terms of both volatile and non-volatile defenses when we considered the geographical distribution of the populations.

## Materials and Methods

### Seed collection and plant cultivation

*Plantago lanceolata* seeds were collected from nine native and ten introduced populations across a wide latitudinal and longitudinal gradient (Fig. 1a, Table S1) throughout the world. In each population, seeds were collected as bulk samples from at least five different plants growing at least 3 m apart. Seeds were cleaned and stored at -20 ° C until the start of the experiment. To sterilized the seeds, we shook them for 2 minutes in a sterilization solution (100 mg sodium dichloro-isocyanurate [DCCA 2% w/v, Sigma, St. Louis, MO, USA] in 5 ml deionized H_2_O + 50 μl Tween 20 [1% v/v, Merck, Darmstadt, Germany] and afterwards rinsed for 30 seconds with tap water. The seeds were then germinated in 0.44 L pots (Ø 10 cm) filled with a mixture of sand and nutrient-poor soil (50:50) under greenhouse conditions in Jena, Germany (20 ± 2 ° C during the day [16 hours)] and 18 ± 2 ° C at night [8 hours], 30 - 55% relative humidity). After germination, 10 seedlings per population were planted individually in pots of the same size and filled with the same substrate as germination pots. Plants were automatically irrigated once a day for 5 minutes and fertilized (0.025% Ferty 3, Planta Düngemittel, Regenstauf, Germany) after 35 days. The arrangement of plants on the greenhouse table was re-randomized once a week to account for variability in light exposure.

**Figure 1.**
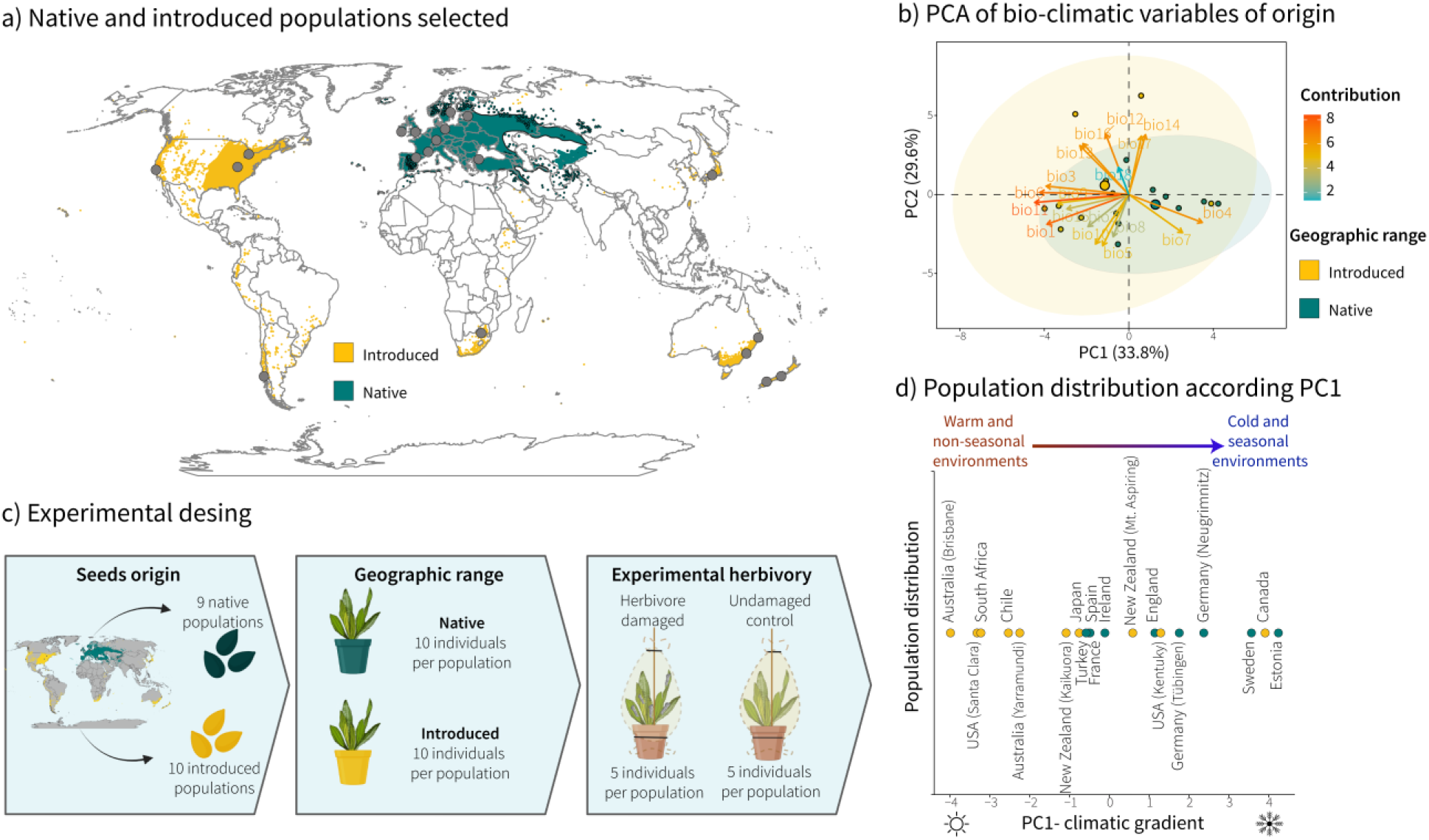
Native and introduced populations of *Plantago lanceolata* selected for this study and experimental design. a) Distribution map of native and introduced ranges of *P. lanceolata*. Dots represent the areas where populations were sampled. b) Principal component analysis results from the first two principal coordinates on the 19 bio-climatic variables extracted from the WORLCLIM data set (Table S2). c) Experimental design. Seeds were collected from nine native and ten introduced populations across all five populated continents. Ten seedlings from each population were germinated and planted in pots under greenhouse conditions. Half of the eight-week-old plants were exposed to five 3rd instar *Spodoptera littoralis* caterpillars for 48 hours, and the other half of the plants functioned as undamaged controls. N = 190 (n = 5 per treatment, per population). d) Populations included in the study and their position along the first principal coordinates of the PCA climatic gradient. Figure created with Rstudio and Biorender.com.

### Environmental variables

Given the wide geographic distribution of the studied populations, we characterized the environment of each sampled population using 19 bio-climatic variables from WorldClim (http://www.worldclim.org). These variables are a set of climatic variables that capture temperature and precipitation annual and seasonal conditions. Variables were extracted on a 2.5-arc min scale based on measurements from 1970 to 2000 (Fick & Hijmans 2017). We conducted a principal components analysis to reduce the axes variation and included the first principal component (hereafter PC1) as an explanatory variable in our model to estimate trait differences between native and introduced populations (see statistical analysis below). PC1 explains 33.3% of the variance among population sites and represents a gradient from sites with warmer temperatures (mean annual temperature and mean temperature of the coldest quarter) and smaller seasonal and day-to-night temperature fluctuations (isothermality), to sites with colder temperatures and larger seasonal and day-night temperature fluctuations (Fig. 1b, Table S2-S3).

### Herbivory treatment

We selected *Spodoptera littoralis* Boisd (Lepidoptera: Noctuidae) as an insect herbivore because caterpillars of this species are broad generalists that will feed on *P. lanceolata*. Caterpillars of *S. littoralis* were hatched from eggs (Syngenta, Basel, Switzerland) and reared in a climate chamber (23± 2°C, with 16 h light per day) on an agar-based artificial diet until they reached the third larval instar. Half of the eight-week-old *P. lanceolata* plants were exposed to five 3^rd^ instar *S. littoralis* caterpillars for 48 hours, and the other half of the plants functioned as undamaged controls (n = 5 per population, per treatment). All plants were enclosed in mesh bags that were tightened at the bottom (at the outer rim of the pots) and at the top with cable ties (Fig. 1c).

### Experimental design

The experiment was staggered over five days, and each day a block of 38 plants (with two plants per population) was processed. To determine chemical defense traits, we first collected volatile organic compounds (VOCs) right after the herbivory treatment. After VOC collection, plant tissues (leaves, inflorescences, root crown and roots) were harvested separately, and immediately flash-frozen in liquid nitrogen and then stored at -80 °C until further chemical analyses. Leaf and root samples were then lyophilized (Alpha 1-4 LD-plus, Martin Christ, Osterode am Harz, Germany) and weighed. Samples were homogenized to a fine powder using a ball mill (MM200, Retsch, Haan, Germany). For chemical analysis, leaf and root powder (10 mg) was extracted with 1 mL methanol for 30 minutes shaking at 240 rpm on a horizontal shaker (IKA® Labortechnik, Steifen im Breisgau, Germany) at room temperature and then centrifuged (2000 g for 5 min) (Method S1).

### Leaf area and leaf damage measurements

To estimate leaf area and experimental leaf damage by *S*.*littoralis* caterpillars, we took pictures of all leaves of each *P. lanceolata* individual right after harvesting. Damage by herbivory was determined by reconstructing the original leaf area with Adobe Photoshop CS5 (Adobe, California, USA; see Unsicker & Mody 2005 for details) to calculate the proportion of total leaf area (cm^2^) and grams consumed (leaf area lost divided by specific leaf area).

### Size-related trait

Number of leaves, leaf area (cm^2^), and plant dry mass (g_dw_), including leaves, roots, flower stem and flower biomass, were analyzed as plant size-related traits taking into account only non-infested plants. Leaf area was calculated as described above.

### Resource-acquisition traits

Specific leaf area (SLA; cm^2^_leaf_ g_dw_^-1^_leaf_), leaf carbon concentrations (mg C g_dw_^-1^) and nitrogen concentrations (mg N g_dw_^-1^) were analyzed as resource-acquisition traits. For analyses of carbon and nitrogen concentrations, approximately 10 mg homogenized leaf material were weighed into tin capsules and measured with an Elemental Analyzer (Vario EL Cube, Elementar, Hanau, Germany).

### Untargeted analysis

Untargeted metabolic profiles for *P. lanceolata* populations leaves were obtained by ultra-high performance liquid chromatography coupled via electrospray ionization (ESI) to a qTOF mass spectrometer (UHPLC-ESI-HRMS), using both the positive and negative ionization modes. The mobile phases consisted of 0.1% v/v formic acid in water and acetonitrile. Raw data files from UHPLC-HRMS were transferred to MetaboScape (Bruker, Germany) to perform bucketing based on MS1 spectra (Method S2). In order to have additional information about the classes of compounds detected in *P. lanceolata we used* SIRIUS (version 5.6.3) classification tool for systematic compound class annotation (Duhrkop *et al*. 2021).

### Targeted chemical analysis

Targeted analyses for leaf and root tissues of iridoid glycosides, verbascoside, flavonoids, and phytohormones were conducted using an HPLC-MS/MS system (HPLC 1260 Infinity II [Agilent Technologies, Santa Clara, USA] - QTrap® 6500+ [AB Sciex, Waltham, Massachusetts, USA]) in multiple reaction monitoring (MRM) mode. For the iridoid glycosides and verbascoside, identification was based on comparison to authentic standards (aucubin: Carl Roth, Germany; catalpol: Wako, Japan; verbascoside: Extrasynthese, France) and quantitative data for these compounds and the phytohormones were calculated by means of internal standards applying experimentally determined response factors when necessary (Method S3, Table S4). For flavonoids (apigenin 7-O-glucoside, luteolin, luteolin-7-glucoside, rutin, quercitrin) relative concentrations were calculated by dividing the peak area by the weight of the sample and the peak area of the internal standard D6-JA.

### VOC analysis

Volatile organic compound (VOC) emissions of *P. lanceolata* were measured using a closed push-pull system for 3 hours. Single plants were enclosed with PET bags that were tightened on top with a cable binder and with a rubber band right at the upper rim of the pots (Fig S1). Compressed air entered the system after passing through an activated charcoal filter (0.7 L/min) and it was pumped out (0.4 L/min) at the top through a Poropak-Q absorbent filter (Volatile Collection Trap (VCT) LLC, USA; Fig. S1). After VOC collection, the traps were eluted with 200 μl dichloromethane containing nonyl acetate (SigmaAldrich, 10 ng μl^-1^), as an internal standard. Subsequently, VOCs were analyzed using a GC-MS chromatograph with helium as the carrier gas for identification and GC-FID for the quantification (Method S4).

## Statistical analyses

To test whether size-related traits, resource-acquisition traits and herbivore damage differ between native and introduced ranges of *P. lanceolata* populations, we performed mixed-effects models. For size-related traits, only undamaged control plants were taken into account, whereas for herbivore damage, we performed generalized mixed models only for herbivore-damaged plants. “PC1-climatic gradient” (continuous variable, Fig 1d) and “range” (factor with two levels) and their interactions were treated as fixed effects and “population” (population nested within range) and “harvest day” as crossed random effects. Investment in reproduction may influence the chemical defense and response to herbivory. Since several individuals of the introduced populations flowered during the experiment, we included flowering of plants (as a factor with two levels for non-flowering vs. flowering) as a covariate that we entered before the other explanatory variables in the model to account for variation attributable to the reproductive stage (y∽ Flowering +PC1* Range + [1|Range/Population] + [1|Harvest date]). We tested the significance of the fixed effects using Type I Wald tests with a chi-square statistic. When needed, data was transformed to meet the assumptions of normality and homogeneity of variances (Table S5-S6).

To test for differences in chemical traits between the native and introduced populations, we performed mixed effect models using the same mixed models as described before but adding “herbivory” (factor with two levels) and their interactions as fixed factors (y∽ Flowering +PC1* Range * Herbivory + [1|Range/Population] + [1|Harvest date]). To test for differences in untargeted metabolite composition between the native and introduced populations, we performed a Partial Least-Squares Discrimination Analysis (PLS-DA) and a heatmap of relative abundances of metabolome features and VOC emission using the same mixed models. We calculated the metabolome diversity via Hill number measures (Chao, Chiu & Jost 2014). In analogy to Hill number measures for species, we used metabolite features and VOC identities, and abundance quantified as peak intensity derived from UHPLC-ESI-HRMS traces and concentration of the VOCs derived from GC-FID analyses (Morris *et al*. 2014). All analyses were run in R version 4.3 (R Development Core Team 2023) using the packages, *raster* and *sp* for worldclim data extraction and map visualization (*Pebesma & Bivand 2005*; *Hijmans 2021*); *lme4, lmerTest, performance, factoextra, mixOmics* and *hillR* for statistical and diversity analysis (Bates *et al*. 2015; Kuznetsova, Brockhoff & Christensen 2017; Rohart et al. 2017; Li 2018; Kassambara & Mundt 2020; Lüdecke *et al*. 2021); *notame* for filtering false positive signal of untargeted metabolites (Klåvus, Kokla et al. 2020); and *ggplot2, pheatmap* and *circlize* for graphical visualization (Gu *et al*. 2014; Wickham 2016).

## Results

### Plant morphological traits and herbivory damage

Several morphological traits (biomass, SLA, proportion of flowering plants) and herbivory damage showed clinal patterns in relation to the original geographical distribution of *P. lanceolata* populations, and the patterns varied between native and introduced populations (Fig. 2). Both leaf and inflorescence biomass were significantly lower in populations from cooler and more seasonal environments (Fig. 2a, Range = *x*^*2*^ *=* 6.25, *P =* 0.012). Introduced populations had more leaves, started flowering earlier, and had a higher proportion of flowering plants (33%) at harvest date, compared to native populations (3%) (Table S6). Specific leaf area (SLA) was higher in populations from cooler and more seasonal environments for native but not introduced populations (Fig 2c, PC1 x Range: *x*^*2*^ *=* 4.09, *P =* 0.042). Plants from introduced populations had lower SLA due to higher leaf mass, rather than smaller leaf area. While foliar nitrogen concentration did not differ among populations, foliar carbon concentration was higher in populations from introduced ranges, and overall, carbon concentrations decreased in populations from cooler and more seasonal environments (Fig. 2d, PC1 = *x*^*2*^ *=*4.52, *P =* 0.034, Range: *x*^*2*^ *=* 7.35, *P =* 0.006).

**Figure 2.**
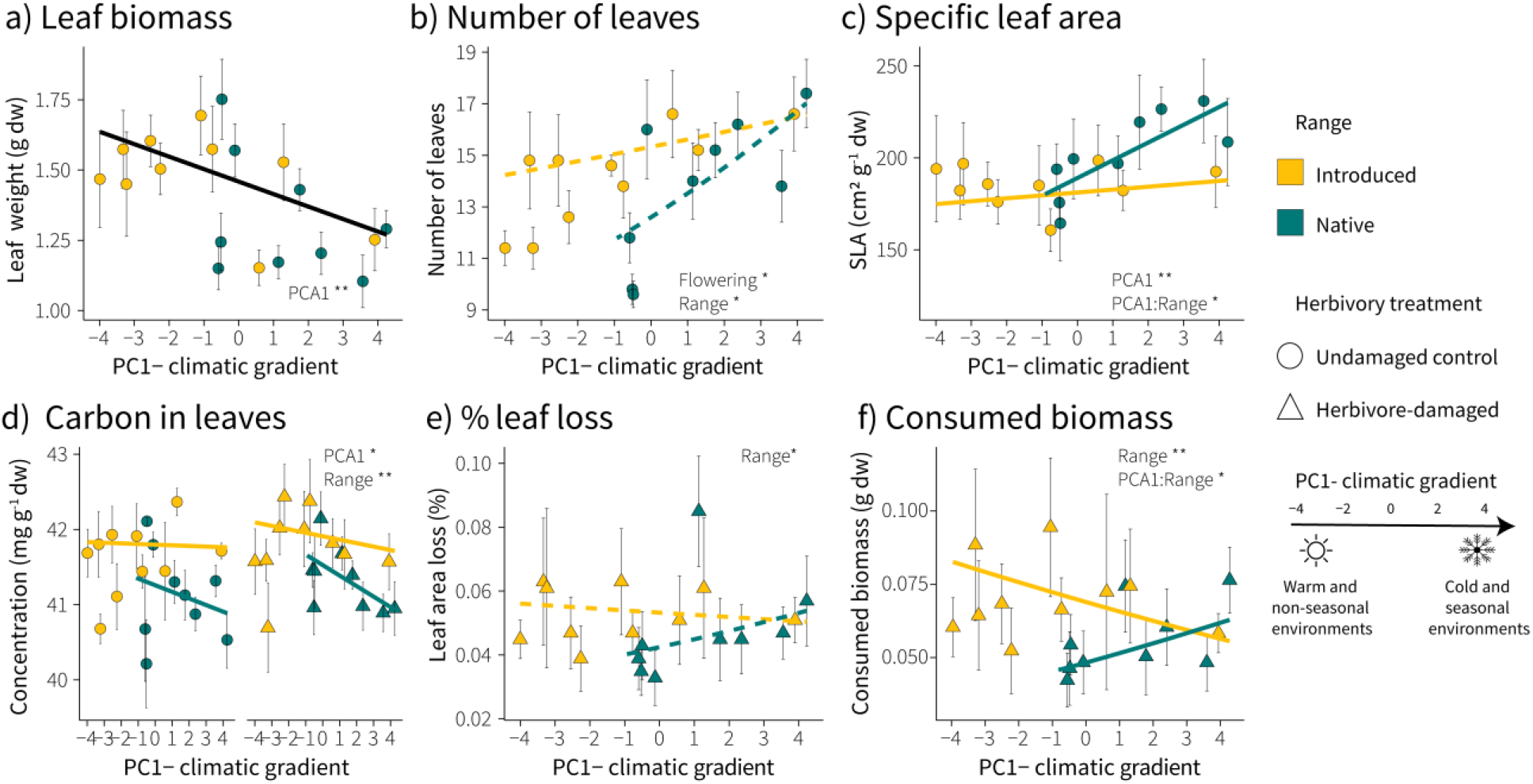
Effect of climatic conditions at population sites (native and introduced populations) on plant morphological traits and herbivore damage in *Plantago lanceolata*. a) Leaf biomass, b) number of leaves, c) specific leaf area (SLA), d) foliar carbon concentration, e) % of leaf loss, and f) consumed biomass by *S. littoralis* caterpillars. Each point (mean ± 1 SE) within a panel represents a single population. Size-related traits were based on undamaged control plants only. For carbon concentration, herbivore damaged plants were included in the model. Broken lines denote non-gnificant relationships with PC1 of the climactic gradient. Asterisks indicate significant effects (*P < 0.05; **P < 0.01; ***P < 0.001).

Experimental herbivory by *S. littoralis* caterpillars for two days caused on average 5% leaf area loss, with introduced populations losing slightly more leaf area than native populations (Fig 2e, 5.2% *vs* 4.6% respectively). Consistent with this, caterpillars consumed more biomass from plants of introduced populations as compared to native populations (Fig. 2f, Range: *x*^*2*^ *=*6.78, *P =* 0.009). Similar to plant resource-acquisition traits, we found climate effects on consumed biomass only in native populations, with leaf damage being higher in populations from cooler and more seasonal environments (Fig 2f, PC1 x Range: *x*^*2*^ *=*5.23, *P =* 0.02).

### Plant chemical traits

The metabolic fingerprinting of leaf methanol extracts with UHPLC-ESI-HRMS in negative ionization mode resulted in a total of 1216 metabolic features. The *in silico* classifications of these metabolic features showed that the metabolic profiles of *P. lanceolata* are dominated by iridoid glycosides, secoiridoid glycosides, megastigmanes, phenylethanoid and phenylpropanoid glycosides, flavonoids, simple phenolics, cinamic acids derivatives, and shikimic acid derivatives (Fig. 3a). The PLS-DA model revealed that the metabolomic profiles of *P. lanceolata* plants clustered together based on population range and herbivore damage. Undamaged plants and herbivore damaged plants showed a strong differentiation, explained by the first component (Fig. 3c). In total, 315 features significantly differed between range and herbivory treatment. The heatmap (169 metabolic features, P < 0.01) showed four broad metabolite clusters (Figure 3b). Cluster II contains metabolic features that are relatively high in native populations in contrast to introduced populations. Cluster II includes features that are relative low in herbivore-damaged plants compared to undamaged control plants. In cluster III we found that metabolic features that are relatively high in herbivore-damaged plants compared to undamaged control plants Cluster IV includes features that are relatively high in introduced populations compared to native populations. When we included climate variables (PC1) in the model, metabolite feature diversity showed different clinal patterns between native and introduced populations. Hill Shannon diversity of metabolites of native populations increased towards cooler environments with greater seasonality, while in introduced populations, climate had no effect on diversity (Fig. 3d; PC1 x Range: *x*^*2*^ *=* 9.39, *P =* 0.002). In both population ranges, metabolite diversity increased after herbivory damage (Fig. 3d; Herbivory: *x*^*2*^ *=*14.58, *P <* 0.001). The patterns of metabolic features from UHPLC-ESI-HRMS analysis in positive ionization mode (Table S11, 4654 metabolic features detected) were quite similar to those in negative ionization mode.

**Figure 3.**
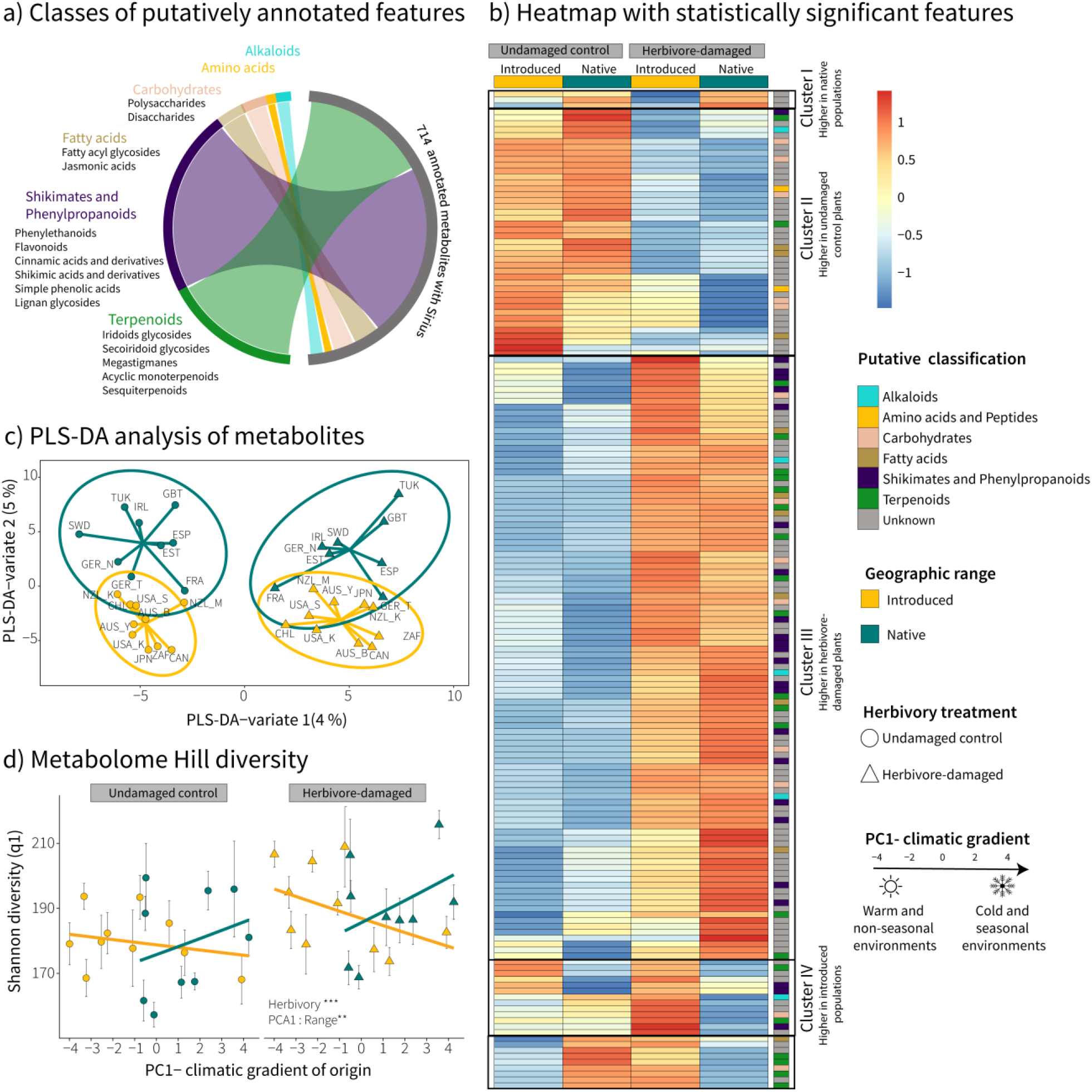
Comparison of native and introduced populations of *Plantago lanceolata* based on untargeted metabolic analysis of methanol extracts. There were 1216 metabolic features in the negative ionization mode after bucketing and filtering. a) Chord diagram showing classes of all the putatively annotated metabolites with SIRIUS, b) Heatmap with most significant features (P < 0.01) affected by population range and herbivory, c) Partial least squares discriminant analysis (PLS-DA) of untargeted metabolites, d) Mean (± 1 SE) of Hill Shannon diversity in relation with PC1-climate gradient. Each point (mean ± 1 SE) within panels’ c and d represents a single population. Prior to analysis, data were normalized by sample weight, log-transformed and Pareto-scaled. Broken lines denote insignificant relationships with PC1 of the climactic gradient. Asterisks indicate significant effects (*P < 0.05; **P < 0.01; ***P < 0.001).

We conducted targeted analysis of leaf and root extracts for iridoid glycosides, verbascoside, flavonoids, and phytohormones using an HPLC-MS/MS system. When we compared native and introduced populations, we did not find differences among plant traits ((Table S7). However, when the climate at the site of origin was taken into account, the foliar concentrations of verbascoside and aucubin were significantly higher in introduced populations, whereas catalpol concentrations did not differ between populations from native and introduced ranges (Fig. 4a-c; aucubin: *x*^*2*^ *=* 7.10, *P =* 0.008; catalpol: *x*^*2*^ *=* 0.52, *P =* 0.476; verbascoside: *x*^*2*^ *=* 7.67, *P =* 0.006). In response to herbivory, verbascoside and iridoid glycoside concentrations in roots and leaves did not increase (Fig. 4a-c, Table S8). Phytohormones in leaves increased after herbivory regardless of population range, with jasmonate concentrations being on average 2.6 times higher and ABA concentrations 1.7 times higher (Fig 4d-c). In general, the relative abundance of flavonoids decreased in populations from cooler, more seasonal environments. After *S. littoralis* exposure, the relative abundance of flavonoids increased in the leaves, primarily due to the increase of luteolin and luteoloside (Fig. 4f, Table S8). Investment in reproduction was associated with changes in the chemical defense of *P. lanceolata*. In general, flowering plants had significantly lower leaf concentrations of verbascoside, aucubin, and the flavonoid quercetin than non-flowering plants (Table S8).

**Figure 4.**
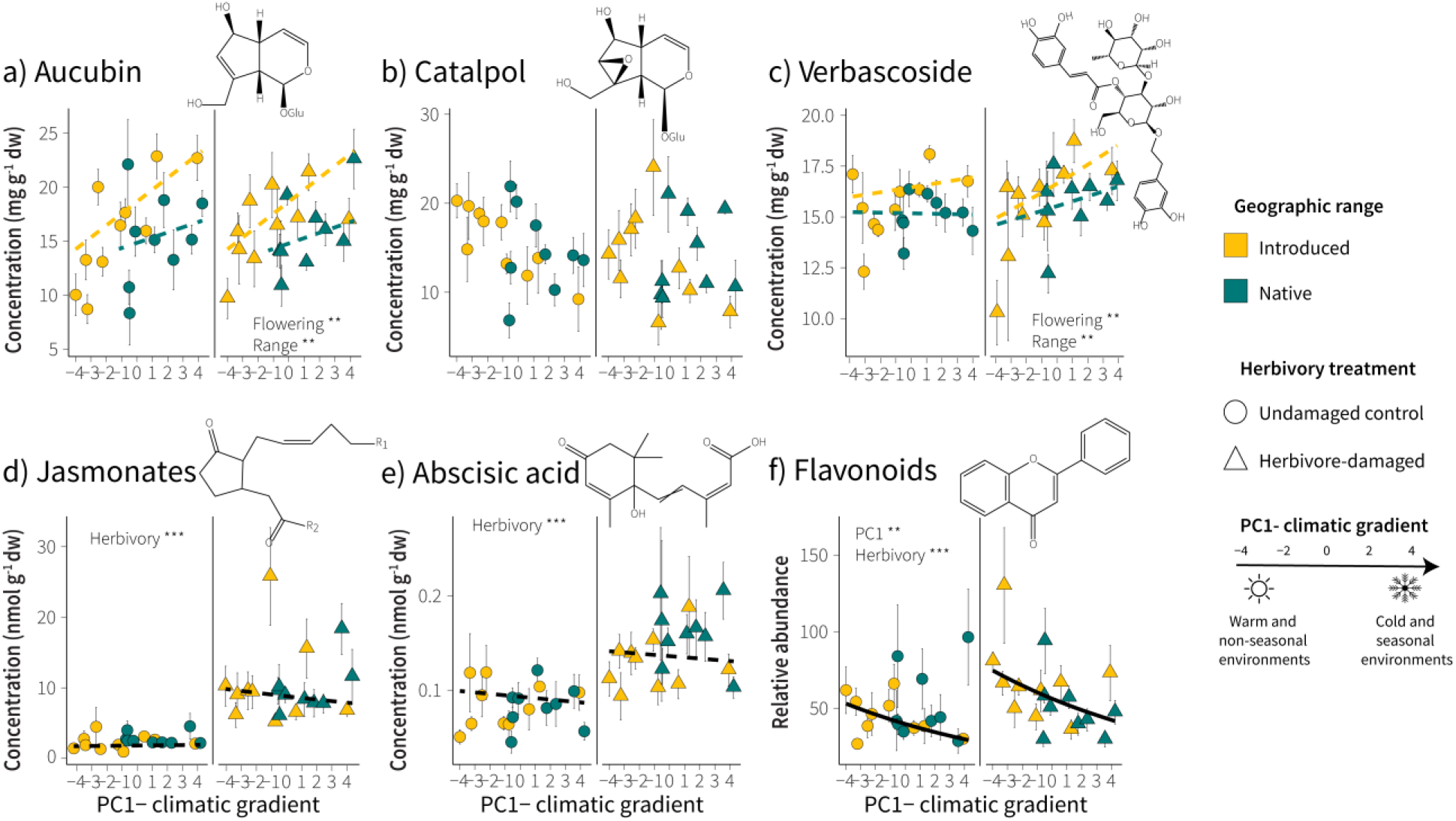
Effect of climatic conditions at population sites (native and introduced populations) and herbivory treatments on levels of characteristic defense compounds and defense hormones in *Plantago lanceolata*. Foliar concentrations of a) aucubin, b) catalpol, c) verbascoside, d) total jasmonates, e) abscisic acid (ABA), and f) total flavonoids of native and introduced populations in relation to PC1 derived from climate variables. Each point (mean ± 1 SE) within a panel represents a single population. Broken lines denote insignificant relationships with PC1 of the climactic gradient. Asterisks indicate significant effects (*P < 0.05; **P < 0.01; ***P < 0.001).

Across the 19 sampled populations, we identified 28 volatile organic compounds (VOCs) released from *P. lanceolata* leaves that belonged to five major groups, namely green leaf volatiles, aromatic compounds, monoterpenes, sesquiterpenes, the homoterpene DMNT and a few other compounds that did not belong to any of these groups (Fig. 5b, Table S9). Terpenes were the largest group of VOCs, but green leaf volatiles were the most abundant. PLS-DA revealed that whereas herbivore-induced VOCs from native and introduced populations of *P. lanceolata* tend to differ, constitutive VOCs emitted from both populations were very similar in their composition. (Fig. 5a-b). Herbivore-induced VOC emission increased in populations from cooler and more seasonal environments, primarily due to the increase of monoterpenes, sesquiterpenes, and green leaf volatiles, with introduced populations showing higher emissions (Fig. 5c, Table S10). After herbivore damage, VOC diversity increased due to the increase of emission and number of compounds. In general, we found differences in VOC profiles of native and introduced populations along climatic gradients. In native populations, VOC diversity decreased toward cooler and more seasonal environments, whereas introduced populations did not show any clinal pattern (Fig. 5d, Table S11).

**Figure 5.**
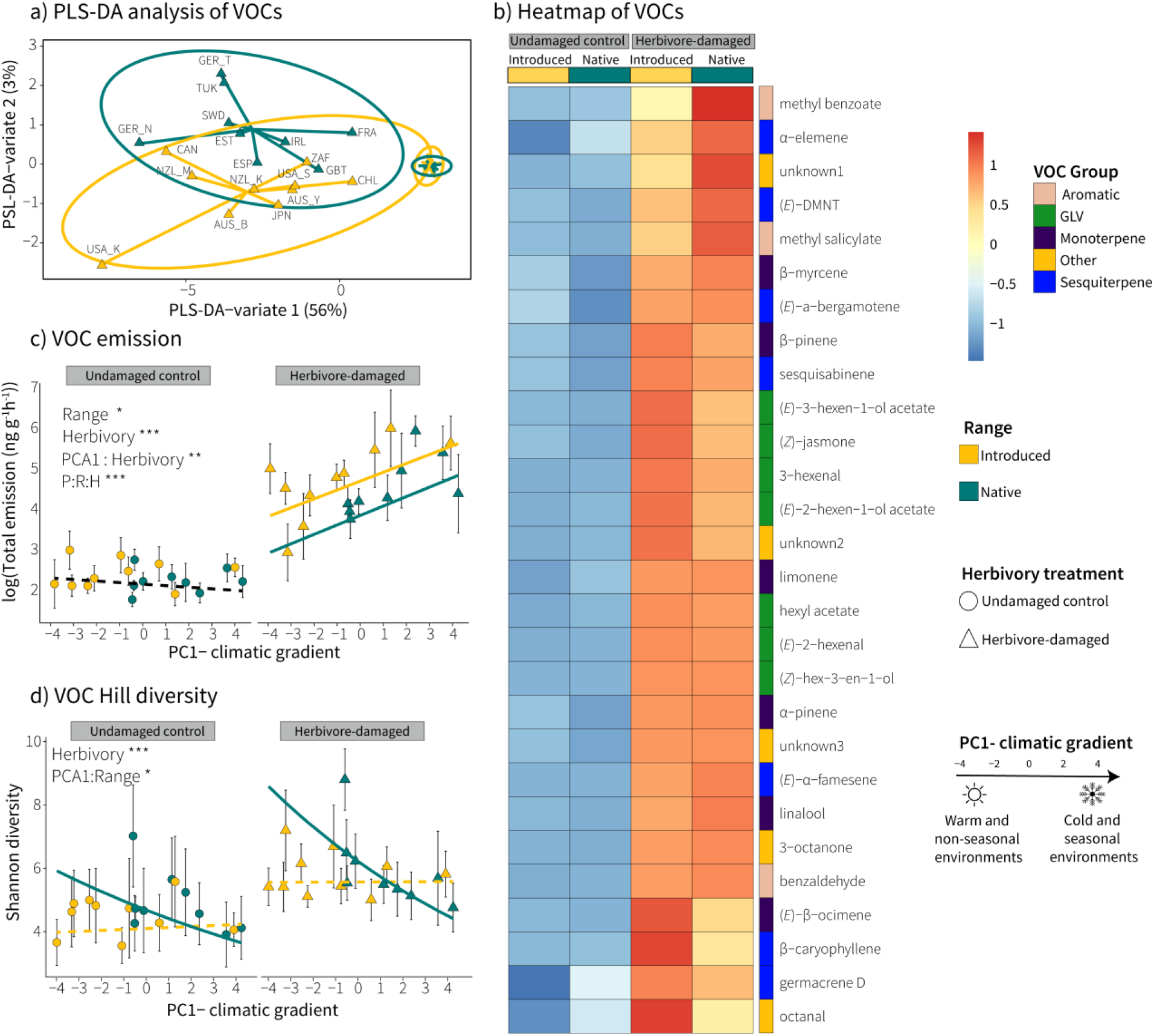
Volatile organic compound (VOC) emission from native and introduced populations of *Plantago lanceolata*. a) Partial least squares discriminant analysis (PLS-DA) of VOCs composition, b) Heatmap of VOCs composition, c) VOC richness in relation with PC1-climate variable, d) VOC Shannon diversity in relation with PC1-climate variable. Each point (mean ± 1 SE) within panels’ c and d represents a single population. Broken lines denote insignificant relationships with PC1 of the climactic gradient. Asterisks indicate significant effects (*P < 0.05; **P < 0.01; ***P < 0.001).

## Discussion

Using a greenhouse experiment, we compared plant size, resource acquisition, and chemical defense of introduced and native populations of *Plantago lanceolata* to investigate whether the establishment of a plant species in a new range leads to shifts in its defense metabolites. Our results showed that when we considered the environmental characteristics of the site of origin, populations from introduced ranges were characterized by an increase of chemical defense compounds without compromising plant biomass, contrary to many of the expectations of the enemy release hypothesis, the evolution of increased competitive ability hypothesis, and the shifting defense hypothesis (Blossey & Nötzold 1995; Keane & Crawley 2002). Our findings illustrate the importance of considering environmental conditions when investigating plant traits and chemical defense divergence between introduced and native populations.

Based on the EICA hypothesis, we expected that introduced populations would invest more in biomass and reproduction. Similar to other studies (Blair & Wolfe 2004; Huang et al. 2010; Villellas et al. 2021), our results were consistent with this trend, with introduced populations starting to flower earlier and having more leaves. Yet, this was mostly because the majority of the introduced populations were from warm and non-seasonal environments, which may select for accelerated plant development (Stamp 2004). In previous work, increases in SLA and leaf nitrogen concentration were also associated with higher growth rates in introduced populations (Leishman et al. 2007). However, introduced *P. lanceolata* populations had lower SLA but similar leaf nitrogen concentrations to those of native populations. Therefore, it remains unclear whether introduced plants invest more in growth in their non-native ranges.

There was little evidence that introduced populations possessed fewer chemical defenses than native populations when we took the climatic conditions of origin into account. Compared to native populations, introduced populations had similar non-volatile metabolic diversity, higher concentrations of aucubin and verbascoside, and higher emission of VOCs. Introduced plants may escape specialist herbivores from their home ranges, but they may still suffer attacks from generalist herbivores in the introduced range (van der Putten *et al*. 2005). Since defense compounds such as iridoid glycosides are more effective on generalists than specialists (Bowers & Puttick 1988; Harvey, van Nouhuys & Biere 2005), the increase in aucubin levels may be considered a logical consequence of the shift from native to introduced ranges. Curiously, we found no induction of iridoid glycosides and verbascoside upon herbivory. These compounds have been reported to be inducible by herbivores in some cases (Fuchs & Bowers 2004), but not all (Stamp & Bowers 1996; Fontana *et al*. 2009), with inducibility depending on the ontogenetic stage of the plant (Quintero & Bowers 2012). In line with the shift in defense strategy toward generalists rather than specialists, we hypothesized that introduced populations would invest in constitutive VOC emission while native populations would invest in herbivore-induced VOCs. However, our findings revealed that introduced populations of *P. lanceolata* emitted higher concentrations of herbivore-induced volatiles compared to native populations, in contrast to earlier studies on *Jacobaea vulgaris* Gaertn. (Lin *et al*. 2021). This suggests that the enemy release hypothesis may not be applicable to all plant species and that different species may have different strategies for dealing with herbivores.

Environmental factors play an important role in structuring plant defense traits, both directly and indirectly, since climate can be a key selective force driving rapid trait adaptations (Colautti, Maron & Barrett 2009; Hahn et al. 2019; van Boheemen et al. 2019). We detected a positive correlation between herbivore-induced VOCs and decreasing temperatures in both native and introduced populations. Similar to our results, studies on *Asclepias syriaca* L. and *Vicia sepium* L. showed a greater induction of VOC emissions at higher latitude/altitude compared to plants at lower latitude/altitude (Wason, Agrawal & Hunter 2013; Rasmann *et al*. 2014). Concentrations of herbivore-induced VOCs have been shown to be positively correlated with the percentage of herbivore damage. However, leaf damage was not correlated with the total emission of herbivore-induced VOCs, suggesting that the increase of VOCs in cooler environments was not explained by herbivore damage. Plant investment in herbivore-induced VOCs at cooler temperatures might reflect a shift to more investment in cost-saving inducible defenses (Rasmann *et al*. 2014) or be a consequence of limited nutrient availability in harsh conditions (Rinnan *et al*. 2014). The shifting composition and concentration of plant metabolites may allow for more robust protection against a range of antagonists and unfavorable abiotic environmental conditions (Shulaev *et al*. 2008). Since the successful establishment of a species depends not only on defense metabolites, other compounds may also play a role in developing adaptive strategies. However, future experiments are needed to test whether these assumptions hold true under field conditions.

Given that introduced plant species are typically introduced into heterogeneous environments containing a variety of levels of biotic and abiotic factors, it is unlikely that all introduced populations will evolve similarly. While we found clinal patterns in some of the measured plant traits, we did not observe consistent responses to climatic variables in all studied traits. In fact, in some cases, responses were even different in native and introduced populations. For example, native populations showed clear clines in SLA and their metabolite diversity, with populations from cooler and more seasonal environments showing leaves with higher SLA and metabolite diversity than native plants in warmer areas, whereas introduced populations showed no variation, a pattern previously reported for SLA in *P. lanceolata* (Alexander *et al*. 2012; Villellas *et al*. 2021). Several traits studied in *P. lanceolata* exhibited clinal patterns in both native and introduced ranges, suggesting rapid adaptation to the local environments following introduction, but this may also result from the joint action of genetic drift and gene flow (Ward, Gaskin & Wilson 2017; Smith *et al*. 2020).

In this study, we integrated targeted and untargeted metabolomics to evaluate how plant growth and defense traits vary between native and introduced *P. lanceolata* populations. Our results now add valuable insights into changes in chemical defense traits in introduced ranges of *P. lanceolata* along climatic gradients. Due to potential maternal effects in our study, environmental-associated clines in plant phytochemical defenses should be interpreted with care. Despite including reproductive status in our models to compensate for plants from warmer areas developing inflorescences earlier than those from cooler populations, such developmental changes can obscure the results (Colautti, Maron & Barrett 2009). It would be interesting to learn more about the underlying mechanisms of the strong correlation between climate and defense levels and why this pattern at a global scale. Follow-up experiments at the chemical and genetic levels under controlled conditions would be helpful in identifying potential drivers. Factors, such as edaphic conditions, plant-plant competition and other biotic variables may be as important as climate and herbivory in influencing chemical defense profiles.

## Acknowledgements

We are grateful to Dylan Child, Patricia Fernandez, Jennifer Firn, Anna Fontana, Almuth Hammerbacher, Robert Junker, Ruth Kelly, Rebeca McCulley, Joslin Moore, Jim Nelson, Miguel Nemesio-Gorriz, Rob Salguero-Gómez and Katja Steinauer for helping us to collect the seed material. We acknowledge the Plantpopnet project coordinated by Yvonne Buckley for support during the collection of seed material. We thank the ICE greenhouse team for growing the plants. Konstantin Albrecht and Petra Hoffmann analyzed carbon and nitrogen concentrations. PMB was supported by the Deutsche Forschungsgemeinschaft (DFG) Research Unit (FOR 5000).

## References

Alexander, J.M., van Kleunen, M., Ghezzi, R. & Edwards, P.J. (2012) Different genetic clines in response to temperature across the native and introduced ranges of a global plant invader. Journal of Ecology, 100, 771–781.

Anstett, D.N., Ahern, J.R., Glinos, J., Nawar, N., Salminen, J.P. & Johnson, M.T. (2015) Can genetically based clines in plant defence explain greater herbivory at higher latitudes? Ecology Letters, 18, 1376–1386.

Bates, D., Mächler, M., Bolker, B. & Walker, S. (2015) Fitting linear mixed-effects models using lme4. Journal of Statistical Software, 67.

Bhattarai, G.P., Meyerson, L.A., Anderson, J., Cummings, D., Allen, W.J. & Cronin, J.T. (2017) Biogeography of a plant invasion: Genetic variation and plasticity in latitudinal clines for traits related to herb. Ecological Monographs, 87, 57–75.

Bischoff, A., Crémieux, L., Smilauerova, M., Lawson, C.S., Mortimer, S.R., Dolezal, J., Lanta, V., Edwards, A.R., Brook, A.J., Macel, M., Leps, J.A.N., Steinger, T. & Müller-Schärer, H. (2006) Detecting local adaptation in widespread grassland species? The importance of scale and local plant community. Journal of Ecology, 94, 1130–1142.

Blossey, B. & Nötzold, R. (1995) Evolution of increased competitive ability in invasive non-indigenous plants: a hypothesis. Journal of Ecology, 83, 887–889.

Bock, D.G., Caseys, C., Cousens, R.D., Hahn, M.A., Heredia, S.M., Hubner, S., Turner, K.G., Whitney, K.D. & Rieseberg, L.H. (2015) What we still don’t know about invasion genetics. Molecular Ecology, 24, 2277–2297.

Bossdorf, O., Auge, H., Lafuma, L., Rogers, W.E., Siemann, E. & Prati, D. (2005) Phenotypic and genetic differentiation between native and introduced plant populations. Oecologia, 144, 1–11.

Bowers, M.D. & Puttick, G.M. (1988) Response of generalist and specialist insects to qualitative allelochemical variation. Journal of Chemical Ecology, 14, 319–334.

Callaway, R.M. & Ridenour, W.M. (2004) Novel weapons invasive success and the evolution of increased competitive ability. Frontiers in Ecology and the Environment, 2, 436–443.

Catford, J.A., Wilson, J.R.U., Pyšek, P., Hulme, P.E. & Duncan, R.P. (2021) Addressing context dependence in ecology. Trends in Ecology & Evolution.

Cavers, P.B., Bassett, I.J. & Crompton, C.W. (1980) The biology of Canadian weeds. 47 Plantago lanceolata Canadian Journal of Plant Science, 60, 1269–1282.

Chao, A., Chiu, C.-H. & Jost, L. (2014) Unifying Species Diversity, Phylogenetic Diversity, Functional Diversity, and Related Similarity and Differentiation Measures Through Hill Numbers. Annual Review of Ecology, Evolution, and Systematics, 45, 297–324.

Cipollini, D., Mbagwu, J., Barto, K., Hillstrom, C. & Enright, S. (2005) Expression of constitutive and inducible chemical defenses in native and invasive populations of Alliaria petiolata. Journal of Chemical Ecology, 31, 1255–1267.

Coley, P.D. & Aide, T.M. (1991) Comparison of herbivory and plant defences in temperate and tropical broad-leaved forests. Plant–Animal Interactions: Evolutionary Ecology in Tropical and Temperate Regions, John Wiley & Sons Ltd., Brisbane, 25–49.

Cremieux, L., Bischoff, A., Smilauerova, M., Lawson, C.S., Mortimer, S.R., Dolezal, J., Lanta, V., Edwards, A.R., Brook, A.J., Tscheulin, T., Macel, M., Leps, J., Muller-Scharer, H. & Steinger, T. (2008) Potential contribution of natural enemies to patterns of local adaptation in plants. New Phytol, 180, 524–533.

Darrow, K. & Bowers, M.D. (1999) Effects of herbivore damage and nutrient level on induction of iridoid glycosides in Plantago lanceolata. Biochemical Systematics and Ecology, 25, 1–11.

Duhrkop, K., Nothias, L.F., Fleischauer, M., Reher, R., Ludwig, M., Hoffmann, M.A., Petras, D., Gerwick, W.H., Rousu, J., Dorrestein, P.C. & Bocker, S. (2021) Systematic classification of unknown metabolites using high-resolution fragmentation mass spectra. Nat Biotechnol, 39, 462–471.

Fick, S.E. & Hijmans, R.J. (2017) WorldClim 2: new 1-km spatial resolution climate surfaces for global land areas. International Journal of Climatology, 37, 4302–4315.

Fontana, A., Reichelt, M., Hempel, S., Gershenzon, J. & Unsicker, S.B. (2009) The effects of arbuscular mycorrhizal fungi on direct and indirect defense metabolites of Plantago lanceolata L. Journal of Chemical Ecology, 35, 833–843.

Fuchs, A. & Bowers, M.D. (2004) Patterns of iridoid glycoside production and induction in Plantago lanceolata and the importance of plant age. Journal of Chemical Ecology, 30, 1723–1741.

Gu, Z., Gu, L., Eils, R., Schlesner, M. & Brors, B. (2014) Circlize implements and enhances circular visualization in R. Bioinformatics, 30, 2811–2812.

Harvey, J.A., van Nouhuys, S. & Biere, A. (2005) Effects of quantitative variation in allelochemicals in Plantago lanceolata on development of a generalist and a specialist herbivore and their endoparasitoids. Journal of Chemical Ecology, 31, 287–302.

Hijmans, R.J. (2021) raster: Geographic Data Analysis and Modeling. R package version 3. 6-20, <https://CRAN.R-project.org/package=raster>.

Huang, W., Siemann, E., Wheeler, G.S., Zou, J., Carrillo, J. & Ding, J. (2010) Resource allocation to defence and growth are driven by different responses to generalist and specialist herbivory in an invasive plant. Journal of Ecology, 98, 1157–1167.

Hunter, M.D. (2016) 8. Linking trophic interactions with ecosystem nutrient dynamics on the phytochemical landscape. The phytochemical landscape. Princeton University Press. 198–251.

Iwanycki Ahlstrand, N., Havskov Reghev, N., Markussen, B., Bruun Hansen, H.C., Eiriksson, F.F., Thorsteinsdottir, M., Ronsted, N. & Barnes, C.J. (2018) Untargeted metabolic profiling reveals geography as the strongest predictor of metabolic phenotypes of a cosmopolitan weed. Ecology & Evolution, 8, 6812–6826.

Kassambara, A. & Mundt, F. (2020) factoextra: Extract and Visualize the Results of Multivariate Data Analyses. CRAN.R-project.

Keane, R.M. & Crawley, M.J. (2002) Exotic plant invasions and the enemy release hypothesis. Trends in Ecology & Evolution, 17, 164–170.

Klåvus A, Kokla M, Noerman S, Koistinen VM, Tuomainen M, Zarei I, Meuronen T, Häkkinen MR, Rummukainen S, Farizah Babu A, Sallinen T, Kärkkäinen O, Paananen J, Broadhurst D, Brunius C, Hanhineva K. (2020) “notame”: Workflow for Non-Targeted LC-MS Metabolic Profiling. Metabolites, 10 (4):135.

Kuznetsova, A., Brockhoff, P.B. & Christensen, R.H.B. (2017) lmerTest Package: Tests in Linear Mixed Effects Models. Journal of Statistical Software, 82.

Li, D. (2018) hillR: taxonomic, functional, and phylogenetic diversity and similarity through Hill Numbers. Journal of Open Source Software, 3, 1041.

Lin, T., Vrieling, K., Laplanche, D., Klinkhamer, P.G.L., Lou, Y., Bekooy, L., Degen, T., Bustos-Segura, C., Turlings, T.C.J. & Desurmont, G.A. (2021) Evolutionary changes in an invasive plant support the defensive role of plant volatiles. Current Biology.

Lüdecke, D., Ben-Shachar, M.S., Patil, I., Waggoner, P. & Makowski, D. (2021) performance: An R Package for Assessment, Comparison and Testing of Statistical Models. Journal of Open Source Software, 6, 3139.

Mack, R.N. & Lonsdale, W.M. (2001) Humans as global plant dispersers: Getting more than we bargained for. Bioscience, 51, 95–102.

Marak, H.B., Biere, A. & Van Damme, J.M.M. (2003) Fitness costs of chemical defense in Plantago lanceolata L.: Effects of nutrient and competition stress. Evolution, 57.

Miehe-Steier, A., Roscher, C., Reichelt, M., Gershenzon, J. & Unsicker, S.B. (2015) Light and nutrient dependent responses in secondary metabolites of Plantago lanceolata offspring are due to phenotypic plasticity in experimental grasslands. PLoS One, 10, e0136073.

Moles, A.T., Wallis, I.R., Foley, W.J., Warton, D.I., Stegen, J.C., Bisigato, A.J., Cella-Pizarro, L., Clark, C.J., Cohen, P.S., Cornwell, W.K., Edwards, W., Ejrnaes, R., Gonzales-Ojeda, T., Graae, B.J., Hay, G., Lumbwe, F.C., Magana-Rodriguez, B., Moore, B.D., Peri, P.L., Poulsen, J.R., Veldtman, R., von Zeipel, H., Andrew, N.R., Boulter, S.L., Borer, E.T., Campon, F.F., Coll, M., Farji-Brener, A.G., De Gabriel, J., Jurado, E., Kyhn, L.A., Low, B., Mulder, C.P.H., Reardon-Smith, K., Rodriguez-Velazquez, J., Seabloom, E.W., Vesk, P.A., van Cauter, A., Waldram, M.S., Zheng, Z., Blendinger, P.G., Enquist, B.J., Facelli, J.M., Knight, T., Majer, J.D., Martinez-Ramos, M., McQuillan, P. & Prior, L.D. (2011) Putting plant resistance traits on the map: a test of the idea that plants are better defended at lower latitudes. New Phytologist, 191, 777–788.

Moreira, X., Castagneyrol, B., Abdala-Roberts, L., Berny-Miery Teran, J.C., Timmermans, B.G.H., Bruun, H.H., Covelo, F., Glauser, G., Rasmann, S. & Tack, A.J.M. (2018) Latitudinal variation in plant chemical defences drives latitudinal patterns of leaf herbivory. Ecography, 41, 1124–1134.

Morris, E.K., Caruso, T., Buscot, F., Fischer, M., Hancock, C., Maier, T.S., Meiners, T., Muller, C., Obermaier, E., Prati, D., Socher, S.A., Sonnemann, I., Waschke, N., Wubet, T., Wurst, S. & Rillig, M.C. (2014) Choosing and using diversity indices: insights for ecological applications from the German Biodiversity Exploratories. Ecology & Evolution, 4, 3514–3524.

Mraja, A., Unsicker, S.B., Reichelt, M., Gershenzon, J. & Roscher, C. (2011) Plant community diversity influences allocation to direct chemical defence in Plantago lanceolata. PLoS One, 6, e28055.

Pebesma, E.J. & Bivand, R.S. (2005) Classes and methods for spatial data in R. R News 5, 5.

Pellissier, L., Roger, A., Bilat, J. & Rasmann, S. (2014) High elevation Plantago lanceolata plants are less resistant to herbivory than their low elevation conspecifics: is it just temperature? Ecography, 37, 950–959.

Penczykowkski, R.M. & Sieg, R.D. (2021) Plantago spp. as models for studying the ecology and evolution of species interactions across environmental gradients. The American Midland Naturalist, 198, 158–176.

Quintero, C. & Bowers, M.D. (2012) Changes in plant chemical defenses and nutritional quality as a function of ontogeny in Plantago lanceolata (Plantaginaceae). Oecologia, 168, 471–481.

R Development Core Team (2023) R: A language and environment for statistical computing. R Foundation for Statistical Computing, Vienna, Austria. Viena, Austria.

Rasmann, S. & Agrawal, A.A. (2011) Latitudinal patterns in plant defense: evolution of cardenolides, their toxicity and induction following herbivory. Ecology Letters, 14, 476–483.

Rasmann, S., Buri, A., Gallot-Lavallée, M., Joaquim, J., Purcell, J., Pellissier, L. & Heard, M. (2014) Differential allocation and deployment of direct and indirect defences by Vicia sepium along elevation gradients. Journal of Ecology, 102, 930–938.

Rinnan, R., Steinke, M., McGenity, T. & Loreto, F. (2014) Plant volatiles in extreme terrestrial and marine environments. Plant Cell Environment, 37, 1776–1789.

Rohart, F., Gautier, B., Singh, A. & Cao, K.-A.L. (2017) mixOmics: An R package for ‘omics feature selection and multiple data integration. PLoS computational biology, 13, e1005752.

Shulaev, V., Cortes, D., Miller, G. & Mittler, R. (2008) Metabolomics for plant stress response. Physiology Plant, 132, 199–208.

Skinner, R.H. & Stewart, A.V. (2014) Narrow-Leaf Plantain (Plantago lanceolata L.) Selection for increased freezing tolerance. Crop Science, 54, 1238–1242.

Smith, A.L., Hodkinson, T.R., Villellas, J., Catford, J.A., Csergo, A.M., Blomberg, S.P., Crone, E.E., Ehrlen, J., Garcia, M.B., Laine, A.L., Roach, D.A., Salguero-Gomez, R., Wardle, G.M., Childs, D.Z., Elderd, B.D., Finn, A., Munne-Bosch, S., Baudraz, M.E.A., Bodis, J., Brearley, F.Q., Bucharova, A., Caruso, C.M., Duncan, R.P., Dwyer, J.M., Gooden, B., Groenteman, R., Hamre, L.N., Helm, A., Kelly, R., Laanisto, L., Lonati, M., Moore, J.L., Morales, M., Olsen, S.L., Partel, M., Petry, W.K., Ramula, S., Rasmussen, P.U., Enri, S.R., Roeder, A., Roscher, C., Saastamoinen, M., Tack, A.J.M., Topper, J.P., Vose, G.E., Wandrag, E.M., Wingler, A. & Buckley, Y.M. (2020) Global gene flow releases invasive plants from environmental constraints on genetic diversity. Proc Natl Acad Sci U S A, 117, 4218–4227.

Stamp, N.E. & Bowers, M.D. (1996) Consequences for plantain chemistry and growth when herbivores are attacked by predators. Ecology, 77, 535–549.

Theoharides, K.A. & Dukes, J.S. (2007) Plant invasion across space and time: factors affecting nonindigenous species success during four stages of invasion. New Phytologist, 176, 256–273.

Unsicker, S.B. & Mody, K. (2005) Influence of tree species and compass bearing on insect folivory of nine common tree species in the West African savanna. Journal of Tropical Ecology, 21, 227–231.

van Boheemen, L.A., Bou-Assi, S., Uesugi, A. & Hodgins, K.A. (2019) Rapid growth and defence evolution following multiple introductions. Ecology & Evolution, 9, 7942–7956.

van der Putten, W.H., Yeates, G.W., Duyts, H., Reis, C.S. & Karssen, G. (2005) Invasive plants and their escape from root herbivory: a worldwide comparison of the root-feeding nematode communities of the dune grass Ammophila arenaria in natural and introduced ranges. Biological Invasions, 7, 733–746.

van Kleunen, M., Bossdorf, O. & Dawson, W. (2018) The ecology and evolution of alien plants. Annual Review of Ecology, Evolution, and Systematics, 49, 25–47.

Villellas, J., Ehrlen, J., Crone, E.E., Csergo, A.M., Garcia, M.B., Laine, A.L., Roach, D.A., Salguero-Gomez, R., Wardle, G.M., Childs, D.Z., Elderd, B.D., Finn, A., Munne-Bosch, S., Bachelot, B., Bodis, J., Bucharova, A., Caruso, C.M., Catford, J.A., Coghill, M., Compagnoni, A., Duncan, R.P., Dwyer, J.M., Ferguson, A., Fraser, L.H., Griffoul, E., Groenteman, R., Hamre, L.N., Helm, A., Kelly, R., Laanisto, L., Lonati, M., Munzbergova, Z., Nuche, P., Olsen, S.L., Oprea, A., Partel, M., Petry, W.K., Ramula, S., Rasmussen, P.U., Enri, S.R., Roeder, A., Roscher, C., Schultz, C., Skarpaas, O., Smith, A.L., Tack, A.J.M., Topper, J.P., Vesk, P.A., Vose, G.E., Wandrag, E., Wingler, A. & Buckley, Y.M. (2021) Phenotypic plasticity masks range-wide genetic differentiation for vegetative but not reproductive traits in a short-lived plant. Ecology Letters, 24, 2378–2393.

Ward, S.M., Gaskin, J.F. & Wilson, L.M. (2017) Ecological genetics of plant Invasion: What do we know? Invasive Plant Science and Management, 1, 98–109.

Wason, E.L., Agrawal, A.A. & Hunter, M.D. (2013) A genetically-based latitudinal cline in the emission of herbivore-induced plant volatile organic compounds. Journal of Chemical Ecology, 39, 1101–1111.

Wickham, H. (2016) ggplot2: Elegant graphics for data analysis. Journal of Statistical Software.

Wolf, V.C., Berger, U., Gassmann, A. & Müller, C. (2011) High chemical diversity of a plant species is accompanied by increased chemical defence in invasive populations. Biological Invasions, 13, 2091–2102.

Yannelli, F.A., Novoa, A., Lorenzo, P., Rodríguez, J. & Le Roux, J.J. (2020) No evidence for novel weapons: biochemical recognition modulates early ontogenetic processes in native species and invasive acacias. Biological Invasions, 22, 549–562.

