## Supplemental Material for "Plant geographic distribution influences chemical defenses in native and introduced *Plantago lanceolata* populations"

**Table S1. *Plantago lanceolata* populations from which seed samples were obtained for this study**. Seeds were collected from at least five different individuals per population as a bulk sample, between 2017 and 2020. To maintain seed viability, seeds were stored at -20°C until the start of the experiment.

| Population | Location | Latitude (DD) | Longitude (DD) | Habitat description |
|---|---|---|---|---|
| **Native populations** | | | | |
| Estonia | Elva | 58.258 | 26.635 | Grassland (grazed) |
| England | Oxford | 51.775 | -1.325 | Woodland |
| France | Avignon | 43.926 | 4.864 | Ruderal grassland |
| Germany | Neugrimnitz | 52.988 | 13.827 | Grassland (grazed) |
| Germany | Tübingen | 48.539 | 9.0371 | Botanical Garden |
| Ireland | Cork-Distillery Fields | 51.900 | -8.486 | Grassland (mown) |
| Spain | Zaragoza | 41.690 | -0.932 | Grassland near river bank |
| Sweden | Uppsala | 59.916 | 17.662 | Urban site |
| Turkey | Istanbul | 41.059 | 28.811 | Ruderal site |
| **Introduced populations** | | | | |
| Australia | Brisbane | -27.505 | 153.077 | Urban forest (open woodland) |
| Australia | Yarramundi | -33.614 | 150.738 | Grassland |
| Canada | Middlesex Centre, Ontario | 43.074 | -81.336 | Rural Area |
| Chile | Tenten | -42.466 | -73.765 | Rural Area (grassland, road) |
| Japan | Kyoto | 34.966 | 135.741 | Urban site |
| New Zealand | Kaikuora | -42.402 | 173.683 | Urban site |
| New Zealand | Mt. Aspiring | -44.509 | 168.744 | Ruderal site |
| South Africa | Pretoria | -25.781 | 28.267 | Urban site |
| United States | Lexington, Kentucky | 38.108 | -84.491 | Grassland |
| United States | Santa Cruz, California | 36.995 | -122.068 | Urban grassland |

**Table S2. Explained variables and loadings of the six principal components from native and introduced populations of *Plantago lanceolata*.** Bioclimatic variables were extracted from WorldClim for each of our 20 study populations on a 2.5-arc min scale based on measurements from 1970 to 2000 (Fick & Hijmans, 2017). Coordinates and contributions (in parenthesis) of the variables. The first PC axis explained 33.8 % of the variation.

| Explained variables | Variable (unit) | PC 1 | PC 2 | PC 3 | PC 4 | PC 5 | PC 6 |
|---|---|---|---|---|---|---|---|
| **bio1** | Mean Annual Temperature (°C) | -0.85 (11.8) | -0.41 (3.2) | 0.26 (2.03) | 0.15 (1.79) | -0.09 (0.86) | -0.07 (0.69) |
| **bio2** | Mean diurnal range (°C) | -0.42 (2.93) | -0.44 (3.71) | 0.4 (4.63) | -0.04 (0.13) | 0.67 (48.28) | 0.08 (0.96) |
| **bio3** | Isothermality (BIO1/BIO7) * 100 | -0.86 (12.04) | 0.12 (0.25) | -0.15 (0.65) | -0.08 (0.53) | 0.46 (22.29) | -0.09 (1.06) |
| **bio4** | Temperature Seasonality (CV) | 0.76 (9.55) | -0.39 (2.82) | 0.46 (6.11) | -0.02 (0.05) | -0.13 (1.67) | 0.21 (5.73) |
| **bio5** | Max Temperature of Warmest Period (°C) | -0.27 (1.2) | -0.71 (9.43) | 0.6 (10.47) | 0.18 (2.46) | -0.07 (0.51) | 0.16 (3.36) |
| **bio6** | Min Temperature of Coldest Period (°C) | -0.92 (13.98) | 0.03 (0.01) | -0.19 (1.07) | 0.2 (3) | -0.23 (5.75) | -0.13 (2.24) |
| **bio7** | Temperature Annual Range ( °C) | 0.56 (5.11) | -0.53 (5.25) | 0.58 (9.95) | -0.03 (0.08) | 0.14 (2.07) | 0.22 (6.39) |
| **bio8** | Mean Temperature of Wettest Quarter (°C) | -0.16 (0.43) | -0.58 (6.42) | 0.57 (9.55) | -0.26 (5.37) | -0.13 (1.78) | -0.41 (23.22) |
| **bio9** | Mean Temperature of Driest Quarter (°C) | -0.73 (8.84) | -0.12 (0.26) | -0.16 (0.76) | 0.52 (21.13) | -0.06 (0.39) | 0.35 (17.03) |

| Explained variables | Variable (unit) | PC 1 | PC 2 | PC 3 | PC 4 | PC 5 | PC 6 |
|---|---|---|---|---|---|---|---|
| **bio10** | Mean Temp. of Warmest Quarter (°C) | -0.34 (1.95) | -0.68 (8.81) | 0.57 (9.5) | 0.15 (1.86) | -0.22 (5.17) | 0.08 (0.94) |
| **bio11** | Mean temperature of coldest quarter (°C) | -0.97 (15.58) | -0.11 (0.23) | -0.05 (0.07) | 0.12 (1.14) | -0.04 (0.17) | -0.14 (2.6) |
| **bio12** | Annual precipitation (mm) | -0.24 (0.95) | 0.84 (13.23) | 0.47 (6.54) | -0.03 (0.05) | -0.04 (0.14) | 0.1 (1.38) |
| **bio13** | Precipitation of wettest period (mm) | -0.51 (4.22) | 0.68 (8.59) | 0.34 (3.41) | -0.3 (7.13) | -0.17 (3.14) | 0.21 (5.9) |
| **bio14** | Precipitation of Driest Period (mm) | 0.17 (0.5) | 0.83 (12.83) | 0.42 (5.06) | 0.3 (7) | 0.12 (1.47) | -0.02 (0.08) |
| **bio15** | Precipitation Seasonality (CV; mm) | -0.64 (6.74) | -0.2 (0.76) | -0.17 (0.89) | -0.64 (32.4) | 0.03 (0.08) | 0.29 (11.5) |
| **bio16** | Precipitation of Wettest Quarter (mm) | -0.48 (3.71) | 0.7 (9.3) | 0.35 (3.62) | -0.29 (6.55) | -0.18 (3.41) | 0.17 (4.06) |
| **bio17** | Precipitation of Driest Quarter (mm) | 0.13 (0.28) | 0.81 (12.25) | 0.44 (5.55) | 0.33 (8.51) | 0.16 (2.81) | 0.02 (0.05) |
| **bio18** | Precipitation of Warmest Quarter (mm) | -0.11 (0.19) | 0.37 (2.63) | 0.83 (20.16) | -0.1 (0.83) | 0.01 (0.01) | -0.31 (12.79) |

**Table S3. Sampled population loading in a principal component analysis (PCA) based on their climatic variables extracted from WorldClim**. Coordinates of each population based on their bioclimatic variables extracted from WorldClim. Variables were extracted on a 2.5-arc min scale based on measurements from 1970 to 2000 (Fick & Hijmans, 2017). Coordinates and contributions (in parenthesis) of the variables. The first PC axis explained 33.8 % of the variation.

| Populations | PC 1 | PC 2 | PC 3 | PC 4 | PC 5 | PC 6 |
|---|---|---|---|---|---|---|
| **Native** | | | | | | |
| Estonia | 4.24 (15.58) | -0.58 (0.33) | -0.55 (0.47) | -1.61 (10.74) | -0.26 (0.37) | 0.05 (0.02) |
| England | 1.14 (1.13) | 0.3 (0.09) | -2.17 (7.22) | 0.82 (2.76) | 0.05 (0.01) | -0.75 (4.05) |
| France | -0.48 (0.2) | -1.84 (3.36) | -0.24 (0.09) | 1.19 (5.9) | -0.16 (0.15) | 0.88 (5.5) |
| Germany (Neugriminitz) | 2.37 (4.86) | -0.86 (0.74) | -0.74 (0.85) | -0.21 (0.19) | -0.35 (0.7) | -0.64 (2.95) |
| Germany (Tübingen) | 1.75 (2.67) | -0.13 (0.02) | -0.17 (0.04) | -0.46 (0.89) | 0.15 (0.13) | -0.71 (3.62) |
| Ireland | -0.11 (0.01) | 2.18 (4.72) | -2.16 (7.13) | 0.82 (2.8) | -0.6 (1.99) | -0.7 (3.49) |
| Spain | -0.51 (0.23) | -3.15 (9.82) | -0.49 (0.37) | 1.6 (10.6) | -0.15 (0.13) | 0.45 (1.43) |
| Sweden | 3.57 (11.01) | -0.45 (0.2) | -1.11 (1.88) | -1.07 (4.74) | -0.24 (0.32) | -0.16 (0.17) |
| Turkey | -0.59 (0.3) | -1.19 (1.4) | -1.03 (1.62) | 0.86 (3.08) | -1.35 (10.21) | 1.6 (18.18) |
| **Introduced** | | | | | | |
| Australia (Brisbane) | -3.99 (13.77) | -0.89 (0.79) | 1.57 (3.77) | 0.23 (0.21) | -0.67 (2.51) | -1.38 (13.49) |
| Australia (Yarramundi) | -2.25 (4.38) | -1.47 (2.13) | 1.19 (2.16) | 0.21 (0.19) | 0.83 (3.87) | -0.88 (5.58) |
| Canada | 3.91 (13.23) | -0.57 (0.32) | 1.69 (4.4) | -0.37 (0.56) | 0.63 (2.19) | 0.02 (0) |
| Chile | -2.53 (5.56) | 5.09 (25.61) | -0.92 (1.3) | -1.23 (6.26) | -1.43 (11.38) | 0.7 (3.47) |
| Japan | -0.76 (0.5) | 0.07 (0) | 4.79 (35.19) | -0.94 (3.67) | -1.76 (17.41) | -0.01 (0) |
| New Zealand (Kaikuora) | -1.09 (1.02) | 0.87 (0.74) | -1.71 (4.46) | 1.3 (6.98) | -0.02 (0) | -1.03 (7.51) |
| New Zealand (Mt. Aspiring) | 0.59 (0.3) | 6.25 (38.68) | 1.8 (4.96) | 0.97 (3.94) | 1.84 (18.98) | 0.31 (0.67) |
| South Africa | -3.23 (9.01) | -2.2 (4.81) | 0.36 (0.2) | -1.63 (11) | 1.53 (13.17) | -0.33 (0.76) |
| USA (Kentucky) | 1.29 (1.44) | -0.69 (0.47) | 2.41 (8.91) | 1.3 (7.01) | 0.78 (3.42) | 1.29 (11.83) |
| USA (Santa Cruz) | -3.32 (9.55) | -0.72 (0.52) | -2.52 (9.72) | -1.78 (13.2) | 1.18 (7.81) | 1.3 (12.02) |

**Method S1: Leaves and roots extraction**

Lyophilized leaf and root powder (10 mg) was extracted with 1 ml methanol. The suspension was homogenized shaking for 30 s in a paint shaker (COMPANY), followed by 30 min shaking at 240 rpm on a horizontal shaker (IKA® Labortechnik, Steifen im Breisgau, Germany) and centrifuged at 3200 rpm for 5 minutes. The following internal standards were added to the methanol solution used for extraction: 40 ng D6-abscisic acid (D6-ABA; Toronto Research Chemicals, Toronto, Canada), 40 ng of D6-jasmonic acid (D6-JA; HPC Standards GmbH, Cunnersdorf, Germany), 40 ng D4-salicylic acid (D6-SA; Santa Cruz Biotechnology, USA), and 8 ng of D6-JA-isoleucine conjugate (D6-JA-Ile; HPC Standards GmbH, Cunnersdorf, Germany).

**Method S2: Untargeted metabolic fingerprinting by UHPLC–ESI–HRMS profiles**

For non-targeted analysis of *P. lanceolata* methanol leaf extracts, ultra-high-performance liquid chromatography–electrospray ionization– high resolution mass spectrometry (UHPLC–ESI–HRMS) was performed with a Dionex Ultimate 3000 series UHPLC (Thermo Scientific) and a Bruker timsToF mass spectrometer (Bruker Daltonik, Bremen, Germany). UHPLC was used applying a reversed-phase Zorbax Eclipse XDB-C18 column (100 mm × 2.1 mm, 1.8 µm, Agilent Technologies, Waldbronn, Germany) with a solvent system of 0.1% formic acid (A) and acetonitrile (B) at a flow rate of 0.3 ml/min. The elution profile was the following: 0 to 0.5 min, 5% B; 0.5 to 11.0 min, 5% to 60% B in A; 11.0 to 11.1 min, 60% to100% B, 11.1 to 12.0 min, 100% B and 12.1 to 15.0 min 5% B. Electrospray ionization (ESI) in negative or positive ionization mode was used for the coupling of LC to MS. The mass spectrometer parameters were set as follows: capillary voltage 4.5 KV or 3.5 KV, end plate offset 500 V, nebulizer pressure 2.8 bar, nitrogen at 280 °C at a flow rate of 8 L/min as drying gas. Acquisition was achieved at 12 Hz with a mass range from *m/z* 50 to 1500. At the beginning of each chromatographic analysis, 10 uL of a sodium formate-isopropanol solution (10 mM solution of NaOH in 50/50 (v/v %) isopropanol water containing 0.2% formic acid) was injected into the dead volume of the sample injection for re-calibration of the mass spectrometer using the expected cluster ion *m/z* values. Samples were randomized and quality control samples were injected every 25 samples. A quality control consisted of a pool of 10 µl aliquots of every sample. A blank injected 3 times at the beginning and end of the run was also included. Peak detection was carried out using Metaboscape software (Bruker Daltonik, Bremen, Germany) with the T-Rex 3D algorithm for qTOF data. For peak detection the following parameters were used: intensity threshold of 1500 with a minimum of 10 spectra, time window from 0.4 to 12 min. Adducts of $[M \pm H]^{\pm}$, $[M \pm Na]^{\pm}$, and $[M \pm K]^{\pm}$ (for positive mode) or $[M\text{-}H]^{-}$, $[M \pm Cl]^{-}$, and $[M \pm COOH]^{-}$ (for negative mode) were grouped as a single bucket if they had an EIC correlation of 0.8 or more. *Feature filters* Minimum number of samples: present in 5 of 190; minimum for recursive feature extraction: present in 5 of 190; group filter: present in at least 80% of at least one group.

After data acquisition and peak alignment, the false-positive signals were filtered by using R package Notadame, using a flag detection mode with quality control limit of 0.75 and group limit of 80%.

Annotation of putative metabolites was performed using the program SIRIUS (Dührkop et al., 2019). The mgf file (containing ion information) exported from Metaboscape was applied to SIRIUS (version 5.6.3) which generated predicted molecular formulas for each feature which were ranked using ZODIAC (Ludwig et al., 2020).The chemical taxonomy of the predicted metabolite structures was obtained by CANOPUS (class and subclass; Dührkop et al., 2021).

**Method S3 Quantification of targeted metabolites**

The extracts from leaves and roots of P. lanceolata were analyzed by high performance liquid chromatography (Agilent 1260 HPLC system, Agilent technologies, Santa Clara, USA) coupled to a mass spectrometer (MS) (API 6500, Sciex, Framingham, USA) and equipped with a Turbospray ion source. One µl of extract was separated on a Zorbax Eclipse XDB-C18 column (4.6 x 50 mm, 1.8 µm, Agilent technologies, Santa Clara, USA). Two solvents formed the mobile phase: formic acid (0.05 %) in ultrapure water as solvent A, and acetonitrile as solvent B. The following gradient was used: 0-0.5 min, 5 % B; 0.5-6.0 min, 5-37.4 % B; 6.0-6.02 min, 37.4-80.0 % B; 6.02-7.5 min, 80.0-100 % B; 7.5-9.5 min, 100 % B and 9.52-12 min, 5 % B. The flow rate was 1.1 ml min-1and the column was kept at 20 ºC. In the MS, the liquid effluent was ionized by electrospray ionization in the negative mode (-4500 eV). The turbo gas temperature was set at 650 ºC. Nebulizing gas was set at 60 psi, curtain gas at 45 psi, heating gas at 60 psi, and collision gas to “medium”.

The mass spectrometer was run in multiple reaction monitoring (MRM) mode (Table S4.) MultiQuant™ 3.0.3 (Sciex, Waltham, Massachusetts, USA). Iridoid glycosides and verbascoside were quantified relative to the internal standards for phytohormones applying experimentally determined response factors (Table S2), while for phytohormones, the respective internal standards were used for quantification. Jasmonate profiling included the measurement of jasmonic acid (JA), 12-hydroxyjasmonic acid (OH-JA), Jasmonic acid- isoleucine (JA-Ile) and 12-hydroxy-jasmonoyl-isoleucine (12-OH-JA-Ile). For flavonoids (apigenin 7-O-glucoside, luteolin, Luteolin-7-glucoside, rutin, quercitrin) relative concentrations were calculated by dividing the peak area by the weight of the sample and the peak area of the internal standard D6-JA.

**Table S4. Details of analysis of phytohormones, iridoid glycosides, verbascoside, and flavonoids by LC-MS/MS [HPLC 1260 (Agilent Technologies)-QTRAP6500 (SCIEX)] in negative ionization mode.**

| Q1 | Q3 | RT (min) | Compound | Internal standart | RF | DP | EP | CE | CXP |
|---|---|---|---|---|---|---|---|---|---|
| 136.93 | 93 | 3.3 | SA | D4-SA | 1.0 | -20 | -8 | -24 | -7 |
| 263 | 153.2 | 3.4 | ABA | D6-ABA | 1.0 | -20 | -12 | -22 | -2 |
| 209.07 | 59 | 3.6 | JA | D6-JA | 1.0 | -20 | -9 | -24 | -2 |
| 322.19 | 130.1 | 3.9 | JA-Ile | D6-JA-Ile | 1.0 | -20 | -4.5 | -30 | -4 |
| 225.1 | 59 | 4.4 | OH-JA | D6-JA | 1.0 | -20 | -9 | -24 | -2 |
| 338.1 | 130.1 | 6.0 | OH-JA-Ile | D6-JA-Ile | 1.0 | -20 | -4,5 | -30 | -4 |
| 140.93 | 97 | 3.3 | D4-SA | | | -20 | -8 | -24 | -7 |
| 269 | 159.2 | 3.4 | D6-ABA | | | -20 | -12 | -22 | -2 |
| 215 | 59 | 3.6 | D6-JA | | | -20 | -9 | -24 | -2 |
| 214 | 59 | 3.6 | D5-JA | | | -20 | -9 | -24 | -2 |
| 328.19 | 130.1 | 3.9 | D6-JA-Ile | | | -50 | -4.5 | -30 | -4 |
| 327.19 | 130.1 | 3.9 | D5-JA-Ile | | | -50 | -4.5 | -30 | -4 |
| 391 | 183 | 1.3 | aucubin (formiate adduct) | D6-JA | 3.67 | -20 | -10 | -18 | -10 |
| 407 | 199 | 0.8 | catalpol (formiate adduct) | D6-JA | 2.95 | -20 | -10 | -18 | -10 |
| 632 | 161 | 5.05 | verbascoside | D6-ABA | 2.11 | -20 | -8 | -46 | -5 |
| 431 | 268 | 5.57 | apigenin-7-O-glucoside | relative quantification | | -20 | -8 | -44 | -5 |
| 285 | 133 | 6.82 | Luteolin | relative quantification | | -20 | -8 | -44 | -5 |
| 447 | 285 | 5.11 | Luteolin-7-glucoside | relative quantification | | -20 | -8 | -40 | -5 |
| 609 | 300 | 4.9 | Rutin | relative quantification | | -20 | -8 | -50 | -5 |
| 447 | 301 | 5.52 | Quercitrin | relative quantification | | -20 | -8 | -32 | -5 |

### Method S4: Push-pull system for volatile organic compounds collection

Half of the eight-week-old *Plantago lanceolata* plants were infested with five 3rd instar *Spodoptera littoralis* caterpillars for 48 hours. Afterwards, volatile organic compounds (VOC) emitted from these plants were trapped for three hours on Poropak Q filters (Alltech Florida, USA) using a push-pull system (Fig. S1). After VOC collections, the traps were eluted with 200µl dichloromethane containing nonyl acetate as an internal standard (concentration: 10 ng per µl).

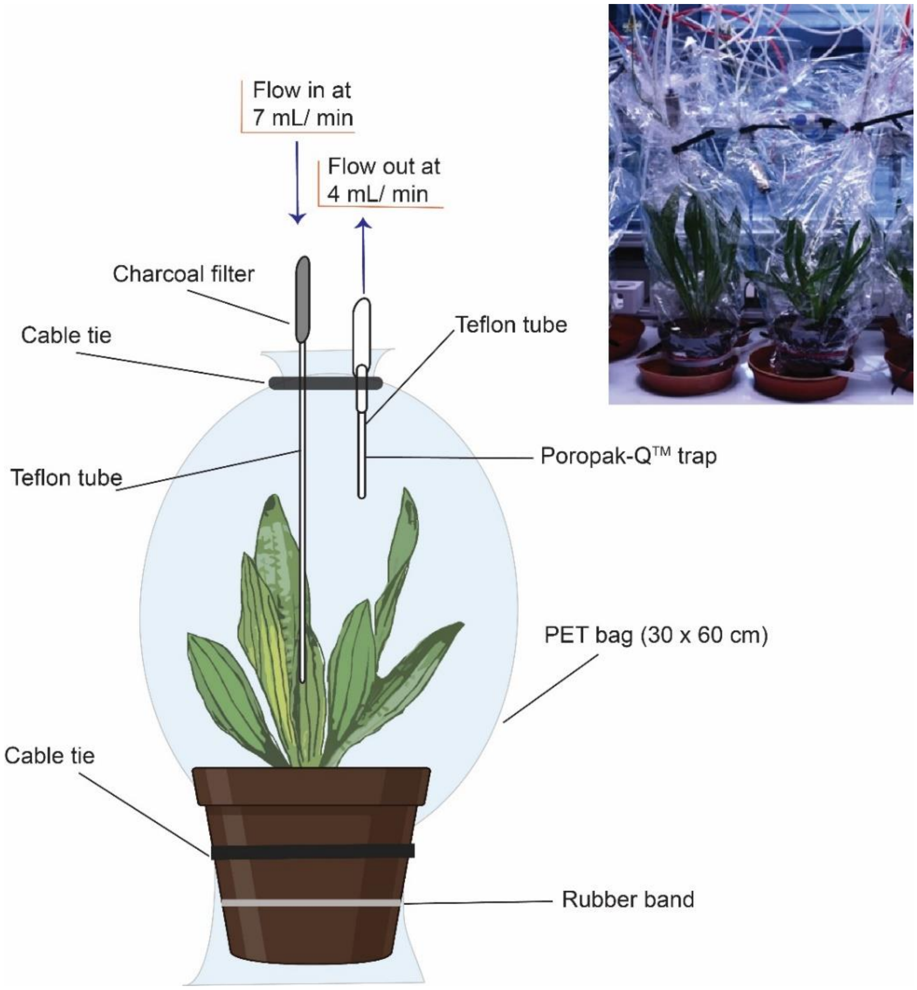


**Figure S1 Scheme of the push- pull system used for VOC collection.** Eight-week-old *Plantago lanceolata* plants were infested with five 3$^{rd}$ instar *Spodoptera littoralis* larvae for 48 hours caged in mesh bags, or were left as undamaged controls. For the volatile collection, the mesh was removed and caterpillars were kept on the plant during the volatile collection. Flowers were excluded from collection bags. Volatile organic compound emissions of *P. lanceolata* were measured for three hours using a push-pull system. Individual plants were enclosed with PET cooking bags (30 cm x 60 cm) and tied on both ends using cable binders (from PET) and rubber bands. Compressed air entered the system after passing through an activated charcoal filter on the lower side of the plant at a flow rate of 7 mL/min and was pumped out at the top through a trap using a vacuum at a rate of 4 mL/min. The trap contained 25 mg Poropak Q adsorbent (Volatile Collection Trap (VCT) LLC, Gainesville, FL, USA) in a Teflon tube that was inserted in the PET bag.

Volatiles were analyzed with a Hewlett-Packard model 6890 gas chromatograph coupled to a Hewlett-Packard model 5973 mass spectrometer (GC-MS). The GC was operated with helium as carrier gas at 1 ml $min^{-1}$, splitless injection (injection temperature: 220 °C, injection volume: 1 µl), a DB-5MS column (30 m × 0.25 mm, 0.25 µm film, J & W Scientific, Folsom, CA, USA), and a temperature program from 40 °C (2 min hold) to 350 °C (2 min hold) with a first gradient of 7 °C $min^{-1}$ to 155 °C, and a second gradient of 60 °C $min^{-1}$ to 300 °C. The GC was coupled to a single quadrupole mass spectrometer operated in electron impact ionization mode. Transfer line temperature was 270 °C, the ionization potential was 70 eV, and a scan range of *m/z* 40–350 was employed.

Volatile compounds were identified by comparing retention times and mass spectra with commercial standards using ChemStation software (Agilent Technologies, Wilmington, DE) and by comparison to the mass spectral libraries Wiley 275 (Wiley and National Institute of Standards) and NIST05a (National Institute of Standards and Technology, Gaithersburg, MD). All standards were purchased from Sigma Aldrich (Saint Louis, MO) (see Table S10). When no commercial standard was available, we used the mass spectrum to identify the compound.

For quantification, the GC was coupled to an FID detector operating at 250 °C. The temperature program had three gradients. Quantification of analytes was accomplished by comparing the peak areas in the FID traces with that of the internal standard calculated and normalized using the following formula:

$$\boldsymbol{E}_{(ng\ \ g^{-1}\,\mathrm{h}^{-1})} = \frac{\boldsymbol{x} \times \boldsymbol{z}_{(ng)} \times \boldsymbol{RF}}{\boldsymbol{y}} \Big/ \boldsymbol{g} \times \boldsymbol{h}$$

Where *E represents the emission in ng per gram per hour*. X and Y are the peak area of the compound and internal standard respectively. Z represents the amount of internal standard used (in this case was 2000 ng) and RF represent the response factor of the compound calculated according to effective carbon number concept of Scanlon and Willis (1985). Concentrations of volatile compounds were standardized for the number of hours (h) for which they were collected and for aboveground plant biomass (g).

**Table S5. Comparison of native and introduced populations of *Plantago lanceolata* based on size-related traits, resource acquisition traits and herbivore damage.** Effects of geographic range (native vs introduced) on size-related and resource- acquisition traits, and herbivore damage were tested using mixed models, including "flowering" and "range" as a fixed effect, and "population" and "harvest day" as random effects. Size-related traits were based on non-infested plants only (45 plant individuals per range). Bold numbers indicate significant effects± P=0.5;*P < 0.05; **P < 0.01; ***P < 0.001). The direction of the arrow indicates either a decrease (↓) or an increase (↑) related to the F (flowering plants), I (introduced populations) or H (herbivore-damaged plants). NA indicates that the factor was not included in the model. Whenever necessary, data were transformed to meet assumptions for linear models.

| Trait | Native | | Introduced | | Model results ($x^2$) | | | | | | |
|---|---|---|---|---|---|---|---|---|---|---|---|
| | Control | Infested | Control | Infested | $R^2$ | Flowering | Range | Herbivory | R:H | F:H | Net effect |
| **Size-related traits** | | | | | | | | | | | |
| Total biomass$^{log}$ | 2.52±0.09 | NA | 2.89±0.09 | NA | 0.09/0.37 | NA | **5.7*** | NA | NA | NA | I↑ |
| Leaves ($g_{dw}$) $^{sqrt}$ | 1.32±0.04 | NA | 1.48±0.04 | NA | 0.07/0.62 | 0.03 | 3.14 | NA | NA | NA | |
| Inflorescences ($g_{dw}$) $^{log}$ | 0 | NA | 0.06±0.01 | NA | 0.4/0.99 | NA | 0.98 | NA | NA | NA | |
| Roots ($g_{dw}$) $^{glmer}$ | 1.19±0.08 | NA | 1.22±0.08 | NA | 0.05/0.26 | 3.69 | 0.44 | NA | NA | NA | |
| R/S ratio | 0.92±0.06 | NA | 0.78±0.07 | NA | | | | | | | |
| Number of leaves $^{glmer}$ | 13.76±0.55 | NA | 14.18±0.44 | NA | 0.11/0.14 | **5.26*** | 2.2 | NA | NA | NA | F↓ |
| **Resource- allocation traits** | | | | | | | | | | | |
| Specific leaf area ($cm^2\ g^{-1}$) $^{sqrt}$ | 201.75±6.57 | NA | 183.59±5.97 | NA | 0.05/0.29 | 0.03 | **4.3*** | NA | NA | NA | I ↓ |
| Foliar Carbon (mg $g_{dw}^{-1}$) | 41.10±0.14 | 41.3±0.11 | 41.61±0.12 | 41.77±0.14 | 0.13/.019 | 1.21 | **9.82**** | 2.15 | NA | 3.58 | I↑ |
| Foliar Nitrogen (mg $g_{dw}^{-1}$) | 1.77±0.03 | 1.76±0.03 | 1.67±0.03 | 1.7±0.02 | 0.05/0.28 | 0.67 | 2.82 | 0.49 | NA | 0.03 | |
| Foliar C:N ratio | 23.40±0.36 | 23.89±0.49 | 25.17±0.35 | 24.57±0.34 | 0.06/0.29 | 0.18 | **3.95*** | 0.069 | NA | 0.0 | I↑ |
| **Herbivore damage** | | | | | | | | | | | |
| Consumed biomass (g) $^{sqrt}$ | NA | 0.055± 0.004 | NA | 0.07 ± 0.006 | 0.06/0.43 | 1.23 | **8.15**** | NA | NA | NA | I↑ |
| Leaf area lost (%) $^{glmer}$ | NA | 0.047± 0.004 | NA | 0.052 ± 0.005 | 0.03/0.40 | 1.17 | 0.03 | NA | NA | NA | |

**Table S6. Effect of climatic conditions at population sites (native and introduced populations) on plant traits and herbivore damage in *Plantago lanceolata*.** Effect of environmental conditions and range (native vs. introduced) on size-related and resource-acquisition traits and herbivore damage were tested using mixed models, including “PC1-climate” and “range” as a fixed effect, and “population” and “harvest day” as random effects. Size-related traits were based on undamaged control plants only. For carbon concentration, herbivore-damaged plants were included in the model. Bold numbers indicate significant effects (*P < 0.05; **P < 0.01; ***P < 0.001). NA indicates that the factor was not included in the model. The direction of the arrow indicates either a decrease (↓) or an increase (↑) related to the factor F (flowering plants), I (introduced populations), N (native populations) or PC1 trend (↗ increase or ↘ decrease). To assess herbivore damage, plants were infested with five 3rd instar *Spodoptera littoralis* caterpillars allowed to feed for 48 hours or left as undamaged controls. When needed, data were transformed to meet statistical assumptions. N=190

| Trait | Model results ($x^2$) | | | | | | | | | |
|---|---|---|---|---|---|---|---|---|---|---|
| | **R2** | **Flowering** | **PC1** | **Range** | **Herbivory** | **P:R** | **P:H** | **R:H** | **P:R:H** | **Net effect** |
| **Size-related traits** | | | | | | | | | | |
| Total biomass | 0.14/0.42 | NA | **4.85*** | 2.58 | NA | 3.37 | NA | NA | NA | ↘ |
| Leaves ($g_{dw}$) | 0.20/0.20 | 0.03 | **7.68**** | 0.7 | NA | 0.01 | NA | NA | NA | ↘ |
| Inflorescences ($g_{dw}$) $^{log}$ | | NA | **12.49 ***** | 0.28 | NA | | NA | NA | NA | ↘ |
| Roots ($g_{dw}$) $^{glmer}$ | 0.1/0.27 | 3.69 | 0 | 0.52 | NA | 3.13 | NA | NA | NA | |
| R/S ratio | 0.21/0.33 | **11.08***** | **5.66*** | 0.13 | NA | 1.74 | NA | NA | NA | F↓ PC1↓ |
| Number of leaves $^{glmer}$ | 0.1/0.27 | **5.26*** | 2.86 | **6.25*** | NA | 3.44 | NA | NA | NA | R↑ |
| **Resource-acquisition traits** | | | | | | | | | | |
| Specific leaf area ($cm^2\ g^{-1}$) | 0.12/0.32 | 0.10 | **8.10**** | 1.50 | NA | **5.03*** | NA | NA | NA | I→ **N↗** |
| Foliar Carbon (mg $g_{dw}^{-1}$) | 0.14/0.18 | 1.21 | **4.52*** | **7.35**** | 2.05 | 1.05 | 0.27 | 0.29 | 0.01 | ↘ I↑ |
| Foliar Nitrogen (mg $g_{dw}^{-1}$) | 0.09/0.28 | 0.67 | 2.27 | 1.46 | 0.51 | 2.8 | 0.48 | 1.13 | 1.29 | |
| Foliar C:N ratio | 0.11/0.29 | 0.21 | 3.18 | 2.31 | 0.13 | 2.98 | 0.39 | 1.78 | 1.33 | |
| **Herbivore damage** | | | | | | | | | | |
| Consumed biomass (g) $^{sqrt}$ | 0.09/0.47 | 1.23 | 1.41 | **6.79**** | NA | **5.24*** | NA | 1.23 | 1.41 | I↑↘N↘ |
| Leaf area lost (%) $^{glmer}$ | 0.04/0.36 | 0.99 | 0 | **3.46*** | NA | 2.39 | NA | 0.99 | 0 | I↑ |

**Table S7. Comparison of native and introduced populations of *Plantago lanceolata* based on levels of defense metabolites and defense hormones.** Effects of population range (native vs. introduced) and herbivory on chemical compounds were tested with mixed-effects model, including flowering as covariate, range and herbivore infestation as a fixed effects, and population and harvest day as random effects. Bold numbers indicate significant effects (*P < 0.05; **P < 0.01; ***P < 0.001). The direction of the arrow indicates either a decrease (↓) or an increase (↑) related to the factor F (flowering plants), H (infested plants), I (introduced populations) and, N (native populations). Plants were infested with five 3rd instar *Spodoptera littoralis* caterpillars allowed to feed for 48 hours or left as undamaged controls. When needed, data were transformed to meet assumptions for linear models.

| Compound | Native (mean ± SE) | | Introduced (mean ± SE) | | Model: variables ($x^2$) | | | | | Net effect |
|---|---|---|---|---|---|---|---|---|---|---|
| | Non-infested | Infested | Non-infested | Infested | Flowering | Range | Herbivory | R:H | F:H | |
| ***Iridoid glycosides** (mg ($g_{dw}$)$^{-1}$)* | | | | | | | | | | |
| Aucubin | | | | | | | | | | |
| *Leaves* $^{sqrt}$ | 15.32±0.94 | 15.77±0.75 | 16.07±0.83 | 16.39±0.86 | **9.27*** | 2.66 | 0.29 | 0.10 | 0.46 | F↓ |
| *Roots* $^{sqrt}$ | 13.53±0.54 | 13.14±0.52 | 12.61±0.46 | 12.96±0.44 | **4.00*** | 0.00 | 0.02 | 0.87 | 0.41 | F↓ |
| *R:S ratio* $^{log}$ | 1.06 ± 0.09 | 1.18 ± 0.13 | 1.02 ± 0.12 | 1.46 ± 0.18 | 3.30 | 2.62 | 0.19 | 0.93 | 0.01 | |
| Catalpol | | | | | | | | | | |
| *Leaves* $^{sqrt}$ | 14.61 ± 0.94 | 14.07 ± 1.02 | 15.77±0.97 | 13.81 ± 0.11 | 0.14 | 0.0 | 2.65 | 0.62 | 2.95 | |
| *Roots* $^{sqrt}$ | 3.77 ± 0.31 | 3.78 ± 0.31 | 3.61±0.28 | 3.62 ± 0.36 | 0.07 | 0.14 | 0.02 | 0.04 | 1.29 | |
| *R:S ratio* $^{log}$ | 0.27 ± 0.02 | 0.32 ± 0.03 | 0.24 ± 0.01 | 0.28 ± 0.02 | 1.81 | 0.32 | 2.43 | 0.10 | 0.38 | |
| Catalpol: Aucubin ratio | | | | | | | | | | |
| *Leaves* $^{cubic}$ | 1.17 ± 0.12 | 0.97 ± 0.08 | 1.24 ± 0.14 | 1.02 ± 0.13 | **4.79*** | **5.26*** | 0.0 | 0.18 | 2.66 | FI↑ |
| *Roots* $^{sqrt}$ | 0.3 ± 0.03 | 0.3 ± 0.03 | 0.3 ± 0.02 | 0.29 ± 0.03 | 0.30 | 3.66 | 0.0 | 0.45 | 2.80 | |
| *R:S ratio log*$^{b}$ | 0.29 ± 0.02 | 0.35 ± 0.03 | 0.3 ± 0.02 | 0.34 ± 0.03 | 2.50 | 0.09 | **5.47*** | 0.17 | 0.03 | H↑ |
| **Caffeoyl phenylethanoid glycoside (mg ($g_{dw}$)$^{-1}$)** | | | | | | | | | | |
| Verbascoside | | | | | | | | | | |
| *Leaves* $^{b}$ | 15.08±0.3 | 15.74±0.42 | 15.67±0.34 | 15.52±0.76 | **7.77**** | 0.01 | 2.75 | 2.52 | **11.07***** | F↓ |
| *Roots* $^{a}$ | 5.18±0.42 | 5.25±0.49 | 4.18±0.29 | 4.14±0.38 | 0.29 | 0.90 | 1.88 | 0.02 | 0.17 | |
| *R:S ratio* $^{b}$ | 0.35 ± 0.03 | 0.35 ± 0.03 | 0.28 ± 0.02 | 0.3 ± 0.02 | 1.69 | 3.20 | 0.23 | 0.48 | 0.78 | |
| ***Flavonoids** (Relative abundance in leaves)* | | | | | | | | | | |
| Apigenin 7-O-glucoside $^{log1}$ | 10.6 ± 1.7 | 10.6 ± 1.72 | 9.93 ± 1.21 | 11.2 ± 1.04 | 0.20 | 0.05 | 0.07 | .59 | 0.36 | |
| Luteolin $^{log1}$ | 10.2 ± 1 | 16.6 ± 2.75 | 9.57 ± 1.36 | 19.4 ± 2.05 | 2.05 | 0.53 | **26.44***** | 1.85 | 2.47 | H↑ |

| Compound | Native (mean ± SE) | | Introduced (mean ± SE) | | Model: variables ($x^2$) | | | | | Net effect |
|---|---|---|---|---|---|---|---|---|---|---|
| | Non-infested | Infested | Non-infested | Infested | Flowering | Range | Herbivory | R:H | F:H | |
| ***Iridoid glycosides** (mg ($g_{dw}$)$^{-1}$)* | | | | | | | | | | |
| Aucubin | | | | | | | | | | |
| Luteolin-7-glucoside [log1] | 11.0 ± 1.87 | 10.2 ± 1.22 | 11.7 ± 1.75 | 16.3 ± 1.70 | 0.06 | 2.89 | **4.72*** | 2.18 | 0.32 | H↑ |
| Rutin [glmer] | 11.9 ± 1.93 | 17.1 ± 6.12 | 13.6 ± 3.43 | 9.81 ± 2.98 | **15.60***** | 0.58 | **7.01**** | **20.67***** | **26.42***** | FH↑ |
| Quercitrin [glmer] | 2.94±0.81 | 2.64±0.84 | 1.84±0.57 | 3.04±1.0 | 0.22 | 0.26 | 3.56 | **13.75**** | **5.77**** | IH↑ FH↑ |
| ***Defense hormones** (ng ($g_{dw}$)$^{-1}$)* | | | | | | | | | | |
| Jasmonic acid (JA) | | | | | | | | | | |
| *Leaves* [a] | 2.13 ± 0.2 | 4.44 ± 0.37 | 1.82 ± 0.17 | 3.73 ± 0.27 | 5.63 | 0.10 | **85.80***** | 0.77 | 0.82 | H↓ |
| *Roots* [a] | 0.87 ± 0.11 | 0.76 ± 0.07 | 0.72 ± 0.08 | 0.73 ± 0.05 | 1.09 | 0.02 | 0.0 | 0.76 | 0.12 | |
| 12-hydroxyjasmonic acid (OH-JA) | | | | | | | | | | |
| *Leaves* [a] | 0.28 ± 0.08 | 2.58 ± 0.34 | 0.17 ± 0.1 | 3.24 ± 0.54 | 0.36 | 0.59 | **154.24***** | 2.91 | 0.24 | H↑ |
| *Roots* | - | - | - | - | - | - | - | - | - | - |
| Jasmonic acid- isoleucine (JA-Ile) | | | | | | | | | | |
| *Leaves* [a] | 0.11 ± 0.01 | 0.15 ± 0.02 | 0.1 ± 0.02 | 0.11 ± 0.01 | **7.79**** | 0.28 | **11.96***** | 0.13 | 0.25 | H↑ |
| *Roots* [a] | 0.05 ± 0.01 | 0.03 ± 0 | 0.04 ± 0.01 | 0.03 ± 0 | 1.36 | 0.98 | **4.80*** | 1.31 | 0.38 | H↓ |
| 12-hydroxy-jasmonoyl-isoleucine (12-OH-JA-Ile) | | | | | | | | | | |
| *Leaves* [a] | 0.01 ± 0 | 0.08 ± 0.01 | 0 ± 0 | 0.08 ± 0.01 | 0.24 | 0.16 | **211.06***** | 0.67 | 0.045 | H↑ |
| *Roots* | - | - | - | - | - | - | - | - | - | - |
| Jasmonates (Total) | | | | | | | | | | |
| *Leaves* [a] | 2.8 ± 0.3 | 9.83 ± 0.91 | 2.26 ± 0.33 | 10.41 ± 1.23 | **4.16*** | 0.21 | **78.69 ***** | 5.82 | 0.55 | H↑ |
| *Roots* [a] | 0.92 ± 0.12 | 0.74 ± 0.07 | 0.75 ± 0.09 | 0.74 ± 0.05 | 2.59 | 0.03 | 0.04 | 0.95 | 0.02 | |
| Abscisic acid (ABA) | | | | | | | | | | |
| *Leaves* [a] | 0.08 ± 0.01 | 0.16 ± 0.01 | 0.09 ± 0.01 | 0.13 ± 0.01 | **5.99*** | 0.28 | **50.83***** | **4.06*** | 1.58 | HN↑ |
| *Roots* | - | - | - | - | - | - | - | - | - | - |



[1] Outliers were detected and samples were removed from the analysis. In three native individuals, relative abundance of luteoloside was 791.67 (Irland), 756.72 (Germany, Neugrimnitz), and 993.41 (Germany, Tubingen). One infested individual had 2046.80 of Luteloside (Canada) and one infested native individual (Germany, Neugriminitz) had 756.72 and 248.27 of luteoloside and apigenin 7-O-glucoside, respectively

**Table S8. Effects of climatic conditions at population collection sites (native and introduced populations) and herbivory treatment on levels of chemical defense compounds in *Plantago lanceolata*.** Results are based on linear mixed-effects models including flowering, environmental conditions, range and herbivore infestation as fixed effects and population and harvest day as random effects. Interactions between fixed effects were included in the model. The table presents $X^2$- values for the fixed effect terms from the Chi-square Wald Test on linear mixed models. Bold values indicate significant differences (× P < 0.06; * P < 0.05; ** P < 0.01; *** P < 0.001). When needed, data were transformed to meet the assumption of normality or homogeneity of variances. The arrows indicate either increased (↑) or reduced (↓) significantly a compound. F: flowering plants, PC: Climate variables loaded in PC1, R: range, H: herbivore infestation. N=190

| **Compound** | | **Model results ($x^2$)** | | | | | | | | |
|---|---|---|---|---|---|---|---|---|---|---|
| | $R^2$ | **Flowering** | **PC** | **Range** | **Herbivory** | **R:H** | **PC:R** | **PC:H** | **PC:R:H** | **Net effect** |
| ***Iridoid glycosides***(mg ($g_{dw}$)$^{-1}$) | 0.04 | **4.92*** | 1.40 | 1.17 | 0.13 | 0.80 | 0.51 | 0.31 | 1.37 | F↓ |
| Aucubin | | | | | | | | | | |
| *Leaves* $^{sqrt}$ | 0.23/0.32 | **9.27**** | 1.86 | **7.11**** | 0.34 | 0.1 | 0.69 | 0.72 | 2.32 | F↓I↑ |
| *Roots* | 0.07/0.28 | **4.01*** | 0.01 | 0.01 | 0.02 | 2.04 | 0.63 | 0.36 | 0.48 | |
| *R:S ratio* $^{log}$ | 0.17/0.24 | 3.3 | 1.53 | **6.69**** | 0.21 | 2.28 | 0.14 | 1.71 | 1.55 | I↓ |
| Catalpol | | | | | | | | | | |
| *Leaves* $^{sqrt}$ | 0.1/0.3 | 0.15 | 2.77 | 0.65 | 2.75 | 3 | 2.15 | 0 | 0.31 | |
| *Roots* $^{sqrt}$ | 0.07/0.37 | 0.07 | 1.87 | 1.34 | 0.03 | 0.52 | 1.19 | 0.18 | 2.9 | |
| *R:S ratio* $^{log}$ | 0.1/0.24 | 1.81 | 0.32 | 0.76 | 2.34 | 1.71 | 0 | 0.14 | **11.34***** | HI↗,HN↘ |
| C:A ratio | | | | | | | | | | |
| *Leaves* $^{cubic}$ | 0.23/0.31 | **5.22*** | **4.44*** | **5.16*** | 3.74 | 1.60 | 2.99 | 0.48 | 0.20 | F↓↘ I↓ |
| *Roots* $^{sqrt}$ | 0.08/0.38 | 0.30 | 3.66 | 1.54 | 0.01 | 0.43 | 0.20 | 1.80 | 2.87 | |
| *R:S ratio* $^{log}$ | 0.14/0.39 | 2.49 | 0.19 | 1.28 | **5.11*** | 0.17 | 0.91 | 0.03 | **9.01*** | HI↗,HN↘ |
| ***Caffeoyl phenylethanoid glycoside*** *(mg ($g_{dw}$)$^{-1}$)* | | | | | | | | | | |
| Verbascoside | | | | | | | | | | |
| *Leaves* $^{glmer}$ | | **17.51***** | 0.81 | **7.67**** | 0.38 | 0.56 | 3.07 | 0.04 | 0.02 | F↓I↑ |
| *Roots* $^{gmler}$ | | 0.06 | 0.63 | 2.04 | 0.57 | 1.51 | 0.32 | 0.41 | **9.8**** | NH↘ |
| *R:S ratio* $^{gmler}$ | | 1.64 | 0.02 | 1.79 | 0.4 | 0.47 | 2.39 | 0.03 | **4.77*** | NH↘ |
| ***Flavonoids*** *(Relative abundance in leaves)* | | | | | | | | | | |

| **Compound** | | **Model results ($x^2$)** | | | | | | | | |
|---|---|---|---|---|---|---|---|---|---|---|
| | $R^2$ | **Flowering** | **PC** | **Range** | **Herbivory** | **R:H** | **PC:R** | **PC:H** | **PC:R:H** | **Net effect** |
| Luteolin $^{log}$ | 0.15/0.25 | 0.18 | 1.21 | 0.046 | **28.27*** ** | 0.86 | 0.08 | 1.93 | 0.60 | H↑ |
| Luteolin-7-glucoside $^{log}$ | 0.12/0.26 | 0.06 | 3.42 | 1.18 | **4.55*** | 0.10 | 0.30 | 1.93 | **6.55*** | CI↘CN↗,HI↑ |
| Rutin $^{glmer}$ | 0.02/0.54 | **15.60*** ** | 0.1 | 0.60 | 11.56 | 0 | **34.12*** ** | **12.84*** ** | 0 | CI↘CN↗,HI↑ |
| Apigenin 7-O-glucoside $^{log}$ | 0.11/0.25 | 0.20 | **6.89** ** | 1.23 | 0.040 | 0.02 | 0.26 | 0.63 | 1.84 | ↘ |
| Quercetin_Rhamnosid $^{glmer}$ | 0.01/0.3 | 0 | 0.16 | 0.16 | **15.69*** ** | 0 | **10.88*** ** | 0.34 | 0.36 | H↑ |
| ***Defense hormones*** *(ng ($g_{dw}$)$^{-1}$)* | | | | | | | | | | |
| Jasmonates (Total) | | | | | | | | | | |
| *Leaves* $^{log}$ | 0.51/0.6 | 2.87 | 0.01 | 0.02 | **157.36*** ** | 0.53 | 0.15 | 2.6 | 2.31 | H ↑ |
| *Roots* $^{sqrt}$ | 0.03/0.27 | 1.46 | 0 | 0.04 | 0.46 | 0.92 | 0.14 | 2.3 | 0.03 | |
| *Root: shoot ratio* $^{sqrt}$ | 0.29/0.52 | 0.67 | 0.06 | 0 | **84.97*** ** | 0.26 | 0.48 | 0.1 | 0.66 | H↑ |
| Jasmonic acid (JA) | | | | | | | | | | |
| *Leaves* $^{sqrt}$ | 0.3/0.5 | 5.98* | 0.54 | 0.06 | **82.72*** ** | 0.32 | 3.11 | 0.56 | 0.95 | H↑ |
| *Roots* $^{glmer}$ | 0.01/0.07 | 1.84 | 0.01 | 0.44 | 1.99 | 0.1 | 0.1 | 1.59 | 0.03 | |
| *R:S ratio* $^{glmer}$ | 0.12/0.3 | 0.43 | 0.03 | 0.06 | **47.38*** ** | 0.02 | 0.02 | 0.03 | 0.04 | H↑ |
| Jasmonoyl isoleucine (JA-Ile ) | | | | | | | | | | |
| *Leaves* $^{sqrt}$ | 0.08/0.12 | 3.45 | 0.15 | 0.81 | 3.7 | 1.37 | **5.27*** | 0.13 | 2.52 | |
| *Roots* $^{glmer}$ | 0.04/0.15 | 0.68 | 0.2 | 0.05 | **6.47*** | 0.07 | 0.19 | 1.46 | 0 | H↑ |
| *R:S ratio* $^{glmer}$ | 0.07/0.2 | 0.02 | 0.33 | 1.76 | **33.62*** ** | 0.28 | 0.06 | **4.05*** | 1.21 | H↑↗ |
| 12-hydroxyjasmonic acid (OH-JA) | | | | | | | | | | |
| *Leaves* | 0.13/0.27 | 0.21 | 0.24 | 0.76 | **92.96*** ** | 1.09 | 0.34 | 3.23 | 0 | H↑ |
| 12-hydroxy-jasmonoyl-isoleucine (12-OH-JA-Ile) | | | | | | | | | | |
| *Leaves* | 0.44/0.49 | 0.31 | 0.32 | 0.11 | **111.87*** ** | 2.19 | 2.34 | 1.18 | **7.45** ** | H↑ |
| Abscisic acid (ABA) | | | | | | | | | | |
| Leaves | 0.75/0.97 | **5.49*** | 0.36 | 0.26 | **48.07*** ** | 0 | 0.62 | 2.85 | 0.54 | H↑ |

**Table S9. Comparison of native and introduced populations of *Plantago lanceolata* based on volatile organic compounds (VOCs) emission. .** Plants were infested with five 3rd instar *Spodoptera littoralis* caterpillars allowed to feed for 48 hours or left as undamaged controls. Compounds are sorted by chemical class and retention time. The emission rate is given in ng $g^{-1}h^{-1}$. Results are based on linear mixed-effects models including flowering, range and herbivore infestation as fixed effects and population and harvest day as random effects. Interactions between fixed effects were included in the model. The table presents $X^2$- values for the fixed effect terms from the Chi-square Wald Test on linear mixed models. Bold values indicate significant differences (× P < 0.06; * P < 0.05; ** P < 0.01; *** P < 0.001). The arrows indicate either increased (↑) or reduced (↓) significantly a compound. F: flowering plants, R: range, H: herbivore-damaged, C: Undamaged control, I: Introduced populations, N: Native populations. † represents compounds identified by comparison to authentic standards, otherwise, compounds were identified by comparison of mass spectra and retention times to those in Willey or Nist Mass library. N=190

| Compound | FID RT | MS RT | Native | | Introduced | | Model $X^2$ | | | | | Net effect |
|---|---|---|---|---|---|---|---|---|---|---|---|---|
| | | | Control | Infested | Control | Infested | Flowering | Range | Herbivory | R:H | F:H | |
| ***Aromatic*** | | | | | | | | | | | | |
| benzaldehyde† | 8.09 | 7.88 | 0.105 ± 0.028 | 0.509 ± 0.067 | 0.096 ± 0.02 | 0.49 ± 0.06 | 0.95 | 0.08 | **98.79*** ** | 0.1 | 0.35 | H↑ |
| methyl benzoate | 11.41 | 11.33 | 0.016 ± 0.009 | 0.344 ± 0.098 | 0.008 ± 0.006 | 0.17 ± 0.07 | 0.05 | 0.87 | **25.11*** ** | 2.32 | 0.2 | H↑ |
| methyl salicylate† | 13.80 | 13.81 | 0.012 ± 0.012 | 0.224 ± 0.074 | 0.02 ± 0.013 | 0.17 ± 0.06 | 0.35 | 0.04 | **16.02*** ** | 0.47 | 1.25 | H↑ |
| ***Green leaf volatiles*** | | | | | | | | | | | | |
| (*E*)-2-hexen-1-ol, acetate | 9.48 | 9.32 | 0.091 ± 0.035 | 1.503 ± 0.301 | 0.072 ± 0.038 | 1.84 ± 0.40 | **11.73*** ** | **5.7*** | **99.27*** ** | 0.14 | 2.19 | F↓I↑H↑ |
| (*E*)-2-hexenal† | 5.66 | 5.2 | 0.035 ± 0.02 | 2.084 ± 0.331 | 0.015 ± 0.007 | 2.14 ± 0.33 | 0.91 | 0.28 | **91.69*** ** | 0.05 | 0.32 | H↑ |
| (*E*)-3-hexen-1-ol acetate† | 9.24 | 9.09 | 10.708 ± 2.873 | 61.263 ± 9.788 | 13.106 ± 2.676 | 76.45 ± 13.92 | 2.05 | **7.52** ** | **110.95*** ** | 0.02 | 0.12 | I↑H↑ |
| (*Z*)-hex-3-en-1-ol† | 5.73 | 5.31 | 1.27 ± 0.451 | 16.572 ± 3.091 | 1.26 ± 0.189 | 16.54 ± 3.35 | 2.19 | 2.15 | **188.02*** ** | 0.02 | 0.54 | H↑ |
| 3-hexenal | 4.49 | 4.15 | 0.302 ± 0.081 | 4.578 ± 0.698 | 0.237 ± 0.031 | 5.23 ± 0.87 | 0.45 | 1.95 | **166.64*** ** | 0.07 | 0.32 | H↑ |
| (*Z*)-jasmone | 18.55 | 18.05 | 0.007 ± 0.005 | 0.046 ± 0.018 | 0.009 ± 0.006 | 0.05 ± 0.02 | 0.01 | 0.08 | **8.45** ** | 0.05 | 0.09 | H↑ |
| hexyl acetate | 9.41 | 9.25 | 0.091 ± 0.051 | 1.196 ± 0.251 | 0.049 ± 0.026 | 1.27 ± 0.29 | **5.81*** | 3.08 | **91.35*** ** | 0.04 | 3.13 | F↓H↑ |
| **Monoterpene** | | | | | | | | | | | | |
| (*E*)- ß -ocimene† | 10.21 | 10.14 | 1.362 ± 0.209 | 10.639 ± 1.327 | 1.581 ± 0.247 | 15.24 ± 2.38 | 0.81 | **4.67*** | **140.34*** ** | 0.48 | 0 | I↑H↑ |
| α -pinene† | 7.40 | 7.19 | 0.025 ± 0.009 | 0.132 ± 0.028 | 0.039 ± 0.009 | 0.129 ± 0.018 | 1.39 | 0 | **43.13*** ** | 0.83 | 1.87 | H↑ |
| ß -myrcene† | 8.79 | 8.67 | 0.224 ± 0.025 | 0.877 ± 0.178 | 0.34 ± 0.026 | 0.805 ± 0.084 | 0.01 | 0.04 | **33.67*** ** | 1.63 | 0.38 | H↑ |

| **Compound** | **FID RT** | **MS RT** | **Native** | | **Introduced** | | **Model $X^2$** | | | | | |
|---|---|---|---|---|---|---|---|---|---|---|---|---|
| | | | **Control** | **Infested** | **Control** | **Infested** | **Flowering** | **Range** | **Herbivory** | **R:H** | **F:H** | **Net effect** |
| ß -pinene† | 8.37 | 8.21 | 0.09 ± 0.02 | 0.55 ± 0.089 | 0.149 ± 0.02 | 0.629 ± 0.077 | 1.09 | 0.36 | **61.46*** ** | 0: | 0.69 | H↑ |
| limonene† | 9.93 | 9.63 | 0.047 ± 0.024 | 0.354 ± 0.092 | 0.01 ± 0.005 | 0.344 ± 0.046 | 0.3 | 0 | **47.22*** ** | 0.02 | 0.06 | H↑ |
| linalool | 11.53 | 11.48 | 0.08 ± 0.013 | 0.423 ± 0.057 | 0.087 ± 0.013 | 0.392 ± 0.051 | 0 | 0.01 | **73.01*** ** | 0.54 | 2.43 | H↑ |
| **Other** | | | | | | | | | | | | |
| 3-octanone† | 8.57 | 8.56 | 0.082 ± 0.018 | 5.316 ± 1.638 | 0.142 ± 0.034 | 5.007 ± 0.609 | 0.23 | 0.02 | **104.51*** ** | 0.63 | 0.05 | H↑ |
| octanal† | 9.13 | 8.99 | 0.171 ± 0.021 | 0.187 ± 0.026 | 0.162 ± 0.02 | 0.207 ± 0.029 | 0.12 | 0.14 | 3.26 | 0.34 | 0.54 | |
| unknown1 | 5.95 | 6.05 | 0.053 ± 0.025 | 1.074 ± 0.242 | 0 | 0.687 ± 0.12 | 3.06 | 0.81 | **66.4*** ** | 2.04 | 0.64 | H↑ |
| unknown2 | 6.01 | 6.23 | 0.019 ± 0.015 | 0.384 ± 0.078 | 0.002 ± 0.002 | 0.485 ± 0.12 | 0 | 0.81 | **49.17*** ** | 0.72 | 0.27 | H↑ |
| unknown3 | 9.00 | 8.83 | 0.009 ± 0.006 | 0.364 ± 0.093 | 0.039 ± 0.014 | 0.352 ± 0.049 | 1.05 | 0.49 | **46.08*** ** | 0.53 | 0.04 | H↑ |
| **Homoterpene** | | | | | | | | | | | | |
| (E)-DMNT† | 11.90 | 11.87 | 0.488 ± 0.124 | 1.886 ± 0.348 | 0.597 ± 0.222 | 1.593 ± 0.277 | **5.3*** | 0.39 | **21.68*** ** | 0.45 | 0 | F↓H↑ |
| **Sesquiterpene** | | | | | | | | | | | | |
| (*E*)- α -famesene | 19.53 | 20.761 | 0.101 ± 0.048 | 1.2 ± 0.254 | 0.128 ± 0.087 | 1.141 ± 0.258 | **4.02*** | 0.29 | **53.32*** ** | 0.01 | 1.92 | F↑H↑ |
| α -elemene | 18.24 | 18.40 | 0.057 ± 0.018 | 0.141 ± 0.052 | 0.025 ± 0.009 | 0.104 ± 0.034 | 0.31 | 0.13 | **9.64**** | 0 | 0 | H↑ |
| ß -caryophyllene† | 18.83 | 19.02 | 0.748 ± 0.194 | 5.271 ± 0.886 | 0.598 ± 0.12 | 8.668 ± 1.989 | 0.19 | **7.18**** | **46.19*** ** | 0.85 | 0.06 | I↑H↑ |
| germacrene D | 20.05 | 20.32 | 0.392 ± 0.187 | 0.584 ± 0.274 | 0.267 ± 0.106 | 0.65 ± 0.155 | 2.8 | 0.03 | **10.1**** | 0.02 | 1.45 | H↑ |
| sesquisabinene | 19.28 | 19.46 | 0.043 ± 0.025 | 0.217 ± 0.07 | 0.067 ± 0.05 | 0.231 ± 0.064 | 0.98 | 0.69 | **11.69*** ** | 0 | 0.24 | H↑ |
| (*E*)- α -bergamotene | 19.08 | 19.32 | 0.052 ± 0.024 | 0.699 ± 0.119 | 0.219 ± 0.129 | 0.68 ± 0.133 | **4.61*** | 3.54 | **31.04*** ** | 1.02 | 0.64 | F↓H↑ |
| **TOTAL** | | | | | | | | | | | | |

**Table S10. Effects of climatic conditions at population collection sites (native and introduced populations) and herbivory treatment on volatile organic compounds (VOCs) emitted by *Plantago lanceolata*.** Plants were infested with five 3rd instar *Spodoptera littoralis* caterpillars allowed to feed for 48 hours or left as undamaged controls. Compounds are sorted by chemical class and retention time. The emission rate is given in ng $g^{-1}h^{-1}$. Results are based on linear mixed-effects models including flowering, environmental conditions, range and herbivore infestation as fixed effects and population and harvest day as random effects. Interactions between fixed effects were included in the model. The table presents $X^2$- values for the fixed effect terms from the Chi-square Wald Test on linear mixed models. Bold values indicate significant differences (× P < 0.06; * P < 0.05; ** P < 0.01; *** P < 0.001). The arrows indicate either increased (↑) or reduced (↓) significantly a compound. F: flowering plants, PC: Climate variables loaded in PC1, R: range, H: herbivore-damaged, C: Undamaged control, I: Introduced populations, N: Native populations. † represents compounds identified by comparison to authentic standards, otherwise, compounds were identified by comparison of mass spectra and retention times to those in Willey or Nist Mass library. N=190

| Compound | FID RT | MS RT | Model $X^2$ | | | | | | | | |
|---|---|---|---|---|---|---|---|---|---|---|---|
| | | | Flowering | PC | Range | Herbivory | P:R | P:H | R:H | P:R:H | Net effect |
| ***Aromatic*** | | | | | | | | | | | |
| benzaldehyde† | 8.09 | 7.88 | 0.95 | 0.52 | 0.37 | **98.88*** ** | 0.75 | 2.88 | 0.38 | 0.14 | H↑ |
| methyl benzoate | 11.41 | 11.33 | 0.05 | 1.26 | 0.33 | **25.39*** ** | 2.89 | 3.53 | 0.79 | 6.41* | H↑ |
| methyl salicylate† | 13.80 | 13.81 | 0.35 | 0.89 | 0.03 | **16.28*** ** | 2.88 | 3.74 | 0.05 | 2.37 | H↑ |
| ***Green leaf volatiles*** | | | | | | | | | | | |
| (*E*)-2-hexen-1-ol, acetate | 9.48 | 9.32 | **11.73*** ** | 2.96 | **8.15** ** | **102.86*** ** | 3 | **4.63*** | 3.67 | 0.22 | F↓ I↑H↑↗C→ |
| (*E*)-2-hexenal† | 5.66 | 5.2 | 0.91 | 2.67 | 1.49 | **92.38*** ** | 0.73 | 1.74 | 0.27 | 0.1 | H↑ |
| (*E*)-3-hexen-1-ol acetate† | 9.24 | 9.09 | 2.05 | 1.06 | **13.75*** ** | **113.44*** ** | 3.19 | 1.04 | 0.1 | 0 | I↑H↑ |
| (*Z*)-hex-3-en-1-ol† | 5.73 | 5.31 | 2.19 | 0.12 | 3 | **189.25*** ** | 2 | 0.01 | 0.13 | 0.28 | H↑ |
| 3-hexenal | 4.49 | 4.15 | 0.45 | 1.26 | **4.4*** | **169.47*** ** | 1.9 | 0 | 0.09 | 0.88 | I↑H↑ |
| (Z)-jasmone | 18.55 | 18.05 | 0.01 | 0.26 | 0.27 | **8.53** ** | 2.59 | 1.47 | 0.81 | 3.84 | H↑ |
| hexyl acetate | 9.41 | 9.25 | **5.81*** | **5.97*** | **6.57*** | **94.56*** ** | 0.78 | **8.63** ** | 3.15 | 0 | F↓I↑H↑↗ C→ |
| **Monoterpene** | | | | | | | | | | | |
| (*E*)- ß -ocimene† | 10.21 | 10.14 | 0.81 | 0.13 | 6.21* | **141.73*** ** | 2.32 | 0.01 | 0.56 | 0.01 | H↑ |
| α -pinene† | 7.40 | 7.19 | 1.39 | 0.26 | 0.08 | **43.00*** ** | 1.39 | **4.45*** | **5.08*** | 2.13 | H↑↗C→<br>IH↑ |

| Compound | FID RT | MS RT | Model $X^2$ | | | | | | | | |
|---|---|---|---|---|---|---|---|---|---|---|---|
| | | | Flowering | PC | Range | Herbivory | P:R | P:H | R:H | P:R:H | Net effect |
| ß -myrcene[†] | 8.79 | 8.67 | 0.01 | 0.1 | 0.01 | **33.66*** ** | 4* | 0 | 1.86 | 1.54 | H↑ |
| ß -pinene[†] | 8.37 | 8.21 | 1.09 | 0.05 | 0.55 | **61.84*** ** | 1.83 | 0.31 | 0.05 | 0.28 | H↑ |
| limonene[†] | 9.93 | 9.63 | 0.3 | 0.07 | 0.02 | **47.36*** ** | 2.26 | 0.04 | 0.01 | **8.39** ** | H↑<br>NH↘ |
| linalool | 11.53 | 11.48 | 0 | 2.95 | 0.43 | **73.88*** ** | 0.61 | **3.89*** | 0.05 | 0.08 | H↑ |
| **Other** | | | | | | | | | | | |
| 3-octanone[†] | 8.57 | 8.56 | 0.23 | 0.04 | 0.05 | **104.63*** ** | **4.22*** | 0.09 | 0.62 | 0.51 | H↑ |
| octanal[†] | 9.13 | 8.99 | 0.12 | 0.12 | 0.31 | 3.31 | 0.73 | 0.4 | 1 | 1.08 | |
| unknown1 | 5.95 | 6.05 | 3.06 | 2.2 | 0.28 | **66.22*** ** | 2.25 | **4.22*** | 0.11 | 0.72 | H↑↗C→ |
| unknown2 | 6.01 | 6.23 | 0 | 0.2 | 1.39 | **49.43*** ** | 0.21 | 0.41 | 1.61 | 0.05 | H↑ |
| unknown3 | 9.00 | 8.80 | 1.05 | 0.35 | 0.27 | **46.06*** ** | 0.31 | 0.25 | 0.31 | 0.73 | H↑ |
| **Homoterpene** | | | | | | | | | | | |
| (*E*)-DMNT[†] | 11.90 | 11.87 | **5.3*** | **11.2*** ** | 2.7 | **33.63*** ** | 3.4 | 3.44 | 0.03 | 0.14 | F↓H↑ |
| **Sesquiterpene** | | | | | | | | | | | |
| (*E*)- α -famesene | 19.53 | 20.761 | **4.02*** | 1.35 | 1.26 | **53.39*** ** | 1.12 | **3.87*** | 0.45 | 0.4 | F↑H↑↗C→ |
| α -elemene | 18.24 | 18.40 | 0.31 | 0.01 | 0.21 | **9.62** ** | 0.21 | 0.21 | 0.11 | 0.07 | H↑ |
| ß -caryophyllene[†] | 18.83 | 19.02 | 0.19 | 1.61 | **5.89*** | **45.87*** ** | 1.02 | 0.03 | 0.71 | 1.01 | H↑ |
| germacrene D | 20.05 | 20.32 | 2.8 | 0.1 | 0 | **10.04** ** | 0.19 | **7.16** ** | **6.04*** | **16.6*** ** | H↑↗C→ |
| sesquisabinene | 19.28 | 19.46 | 0.98 | 0.04 | 0.66 | **11.72*** ** | 0.44 | 0.3 | 0.06 | 0.1 | H↑ |
| (*E*)- α -bergamotene | 19.08 | 19.32 | **4.61*** | 2.4 | **8.52** ** | **33.00*** ** | 1.04 | **9.15** ** | 0 | 0.46 | F↓H↑ |
| **TOTAL[cubic]** | | | 2.17 | 1.20 | **3.95*** | **123.77*** ** | 0.42 | **5.65*** | 2.12 | 0.37 | I↑H↑↗<br>C→ |

**Table S11. Hill number diversity of volatile organic compounds (VOCs) and untargeted metabolites of native and introduced populations of *Plantago lanceolata*.** We calculated metabolome Hill richness and Hill Shannon diversity for each population. Metabolites were used as surrogates for species in the volatile and metabolite diversity data. Concentrations of VOC emission and peak intensity of the non-volatile metabolites from the GC-FID and UHPLC-ESI-HRMS were used as abundance. Plants were infested with five 3rd instar *Spodoptera littoralis* caterpillars allowed to feed for 48 hours or left as undamaged controls. N=190

| Population | Volatile organic compounds (VOCs) | | | | Non-volatile untargeted metabolites: Negative mode metabolites | | | | Non-volatile untargeted metabolites: Positive mode metabolites | | | |
|---|---|---|---|---|---|---|---|---|---|---|---|---|
| | Richness | | Shannon | | Richness | | Shannon | | Richness | | Shannon | |
| | Control | Infested | Control | Infested | Control | Infested | Control | Infested | Control | Infested | Control | Infested |
| **Native** | | | | | | | | | | | | |
| Estonia | 9.6 ± 1.29 | 16.2 ± 3.60 | 3.73 ± 0.86 | 4.76 ± 0.77 | 1096 ± 7 | 1070 ± 27 | 181 ± 6 | 192 ± 5 | 4178 ± 26 | 4129 ± 67 | 796 ± 15 | 803 ± 48 |
| England | 10.4 ± 1.568 | 19.8 ± 1.855 | 5.65 ± 1.31 | 5.49 ± 0.60 | 1109 ± 6 | 1107 ± 14 | 167 ± 5 | 187 ± 9 | 4215 ± 23 | 4206 ± 23 | 820 ± 14 | 827 ± 10 |
| France | 14.5 ± 1.71 | 18.5 ± 2.53 | 4.73 ± 0.37 | 5.54 ± 0.54 | 1108 ± 11 | 1101 ± 9 | 199 ± 11 | 194 ± 5 | 4159 ± 32 | 4150 ± 17 | 846 ± 14 | 831 ± 10 |
| Germany (N) | 8 ± 0.77 | 22 ± 1.47 | 4.57 ± 0.98 | 5.13 ± 0.75 | 1089 ± 3 | 1105 ± 10 | 195 ± 6 | 186 ± 8 | 4211 ± 42 | 4203 ± 23 | 838 ± 18 | 832 ± 29 |
| Germany (T) | 11.2 ± 2.06 | 20.8 ± 3.18 | 5.24 ± 1.37 | 5.34 ± 0.85 | 1098 ± 6 | 1119 ± 11 | 167 ± 3 | 186 ± 7 | 4190 ± 28 | 4295 ± 10 | 805 ± 31 | 904 ± 24 |
| Ireland | 11.25 ± 1.49 | 19.2 ± 1.83 | 4.66 ± 1.36 | 6.22 ± 0.86 | 1083 ± 14 | 1095 ± 11 | 157 ± 4 | 169 ± 4 | 4130 ± 18 | 4147 ± 36 | 786 ± 6 | 836 ± 33 |
| Spain | 10.2 ± 1.02 | 20.4 ± 2.38 | 4.26 ± 0.88 | 6.49 ± 1.04 | 1085 ± 11 | 1084 ± 7 | 188 ± 5 | 206 ± 11 | 4061 ± 41 | 4088 ± 10 | 776 ± 20 | 757 ± 29 |
| Sweden | 9.4 ± 2.66 | 20.8 ± 1.59 | 3.91 ± 1.02 | 5.69 ± 1.49 | 1118 ± 12 | 1114 ± 14 | 196 ± 15 | 216 ± 4 | 4209 ± 47 | 4200 ± 38 | 826 ± 26 | 836 ± 26 |
| Turkey | 10.8 ± 2.60 | 21 ± 1.643 | 7.02 ± 1.62 | 8.8 ± 0.97 | 1069 ± 7 | 1083 ± 10 | 162 ± 6 | 172 ± 5 | 4066 ± 11 | 4107 ± 34 | 761 ± 13 | 843 ± 33 |
| **Introduced** | | | | | | | | | | | | |
| Australia (B) | 8.2 ± 1.85 | 20.4 ± 1.03 | 3.66 ± 0.73 | 5.41 ± 0.59 | 1075 ± 22 | 1112 ± 8 | 179 ± 6 | 207 ± 4 | 4144 ± 37 | 4203 ± 20 | 799 ± 17 | 825 ± 30 |
| Australia (Y) | 11.8 ± 1.53 | 20.4 ± 1.81 | 4.82 ± 1.17 | 5.11 ± 0.34 | 1106 ± 8 | 1094 ± 8 | 182 ± 6 | 204 ± 3 | 4205 ± 18 | 4176 ± 41 | 781 ± 41 | 854 ± 27 |
| Canada | 11.8 ± 0.92 | 21.8 ± 1.39 | 4.06 ± 0.53 | 5.82 ± 0.72 | 1079 ± 3 | 1089 ± 5 | 168 ± 8 | 183 ± 5 | 4124 ± 21 | 4182 ± 10 | 777 ± 28 | 828 ± 37 |
| Chile | 12.25 ± 3.09 | 16.4 ± 1.81 | 4.99 ± 0.97 | 6.15 ± 0.62 | 1086 ± 7 | 1097 ± 7 | 180 ± 8 | 179 ± 9 | 4113 ± 23 | 4183 ± 28 | 796 ± 9 | 831 ± 21 |
| Japan | 11.4 ± 1.86 | 23 ± 0.91 | 4.74 ± 1.57 | 5.44 ± 0.56 | 1085 ± 13 | 1099 ± 8 | 193 ± 7 | 209 ± 12 | 4127 ± 21 | 4195 ± 23 | 800 ± 16 | 850 ± 31 |
| New Zealand | 12.5 ± 0.96 | 22.75 ± 1.25 | 3.56 ± 0.58 | 6.69 ± 1.31 | 1104 ± 12 | 1095 ± 5 | 178 ± 12 | 192 ± 4 | 4181 ± 23 | 4181 ± 29 | 808 ± 34 | 825 ± 19 |
| New Zealand (MA) | 10.4 ± 1.81 | 18.8 ± 2.50 | 4.278 ± 0.90 | 5.01 ± 0.66 | 1097 ± 12 | 1090 ± 11 | 185 ± 7 | 177 ± 7 | 4148 ± 32 | 4159 ± 42 | 801 ± 27 | 787 ± 41 |
| South Africa | 11.25 ± 2.66 | 16.2 ± 4.08 | 4.889 ± 1.05 | 7.197 ± 1.27 | 1086 ± 9 | 1072 ± 8 | 169 ± 6 | 183 ± 6 | 4134 ± 20 | 4071 ± 18 | 810 ± 15 | 753 ± 27 |
| USA (Kentucky) | 9.25 ± 1.03 | 24.2 ± 1.11 | 5.579 ± 1.43 | 6.06 ± 0.62 | 1098 ± 5 | 1098 ± 6 | 176 ± 7 | 174 ± 4 | 4196 ± 4 | 4210 ± 13 | 828 ± 9 | 837 ± 13 |
| USA (Santa Cruz) | 14.8 ± 2.31 | 20.8 ± 1.24 | 4.629 ± 1.11 | 5.41 ± 0.78 | 1094 ± 11 | 1113 ± 9 | 194 ± 4 | 195 ± 5 | 4172 ± 41 | 4178 ± 32 | 808 ± 16 | 877 ± 21 |
